# Maternal motivation overcomes innate fear via prefrontal switching dynamics

**DOI:** 10.1101/2025.01.17.633494

**Authors:** Yunyao Xie, Yijia Li, Xinke Du, Longwen Huang

## Abstract

Parental care is altruistic. In natural environments, parents are often faced with challenging environmental conditions, such as severe weather, complex terrain and predatory threats, and therefore need to overcome the fear of adverse conditions to protect and raise the offspring. Although a few studies have reported risk-taking maternal behaviors^1–3^, it is unknown how maternal motivation and environmental threats are represented and integrated in neural circuits to resolve the conflict and dynamically drive behaviors. Here we report a novel risk-taking maternal behavior paradigm in a semi-naturalistic context, in which a female mouse has to overcome fear and jump off an elevated platform to retrieve pups outside a nest on the ground. We show that while fear of heights reduces the motivation to jump, the presence of pups dramatically facilitates overcoming such fear. A medial prefrontal-periaqueductal gray (mPFC-PAG) pathway is specifically required for the effect of pups on overcoming fear of height, and this circuit integrates conflicting cues about pup and height and encodes motivation to drive risk-taking jumping behaviors. In contrast to cued, fast and predictable reaction timing in typical structured tasks^4,5^, behaviors in our paradigm are highly spontaneous, characterized by stochastic transitions between low-motivation and high-motivation states. Our data reveal that such spontaneity is shaped by the switching ramping dynamics of neural activity in the motivation-encoding dimension, rather than continuous ramping dynamics. Pup and height cues modulate the switching ramping dynamics to influence, but not immediately evoke behaviors. Together, we propose that the prefrontal-brainstem pathway plays vital roles in encoding altruistic motivation to overcome innate fear, and the switching ramping dynamics might represent a general mechanism that gives rise to spontaneous behaviors in naturalistic and conflicting conditions.

## Main

Parental care is indispensable for offspring survival and health. However, complex environmental conditions in habitats, including severe weather, complex terrain and predatory threats, pose a major challenge to parental care: parents, especially mothers, have to overcome their fear of dangers to protect the offspring at the cost of their own safety. The behavioral strategies to resolve the conflict and the underlying neural mechanisms are crucial in understanding animal sociality and altruism. Despite growing knowledge about neural circuits that modulate maternal behaviors^1,6–17^, little is known on how maternal motivation and environmental threats are represented and integrated to dynamically drive risk-taking maternal behaviors.

### Maternal motivation drives risk-taking behaviors

To answer this question, we first developed a novel risk-taking maternal behavior paradigm in mice. We took advantage of experienced virgin mice, as they have been shown to learn maternal behaviors, including pup retrieval, through co-housing with dams and pups and exhibit maternal motivation^8,14^. To introduce conflicts between an environmental threat and pup-directed motivation, we placed an experienced virgin mouse on an elevated platform and scattered pups outside the nest on the ground (Fig. 1a). Fear of height typically results in higher sympathetic activity, prolonged immobility and reduced exploratory behaviors on the elevated platform, and longer latency to jump onto the ground, particularly when the height is greater than 10 cm^18,19^. However, we expected that when pups are present, maternal motivation will facilitate overcoming fear of height and drive female mice to jump off the platform to take care of pups.

**Figure 1.**
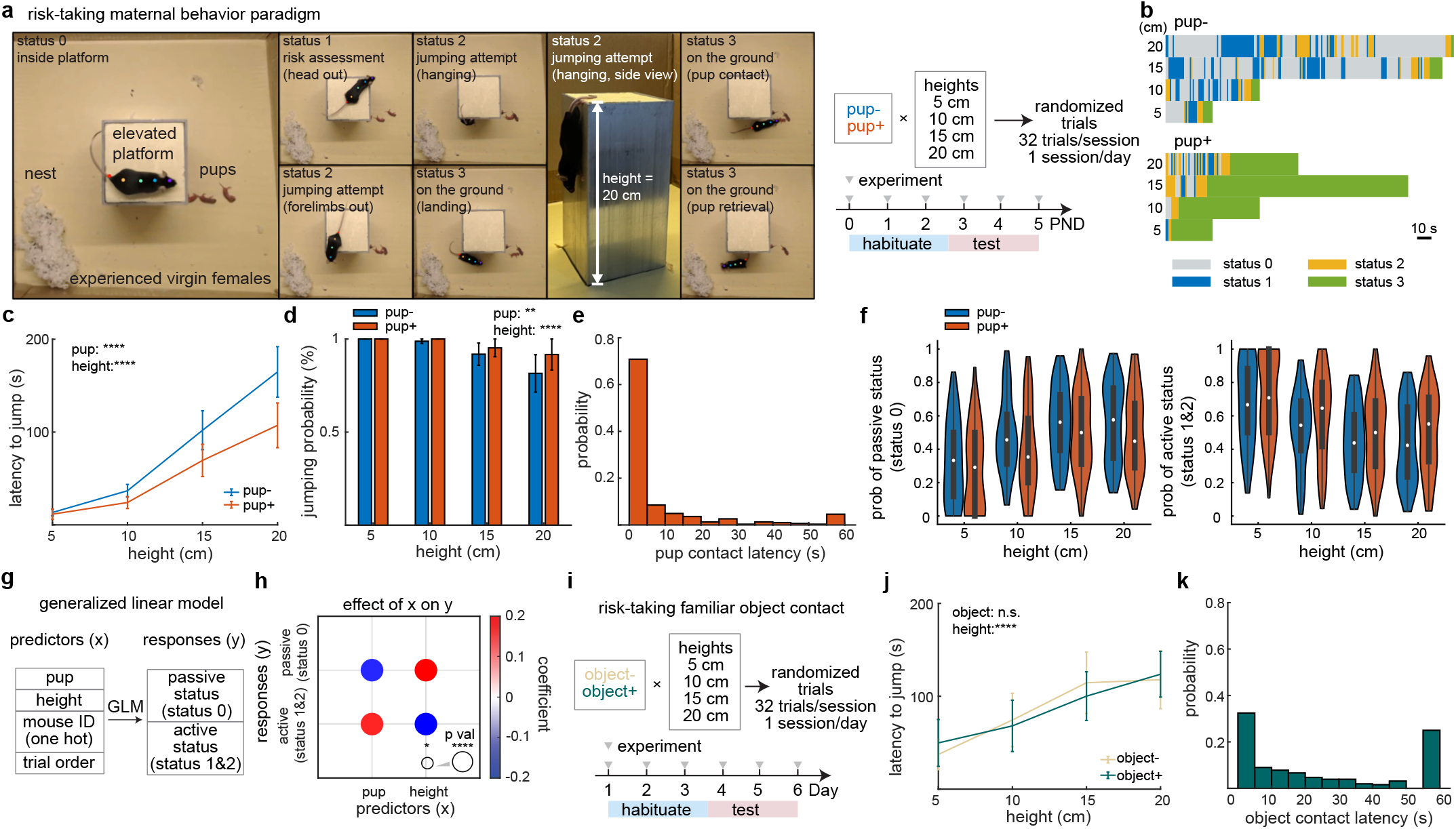
Maternal motivation drives pup-oriented risk-taking behaviors. **a**. Illustration of the risk-taking maternal behavior paradigm. Left, behavioral statuses with example frames of an experienced virgin mouse. Color dots are body parts labeled by DeepLabCut. Right, schematic of the behavior paradigm across post-natal day (PND) 0-5. Each day consists of one session. **b**. Behavioral status series of example trials within the same session. **c**. Latency to jump off the elevated platform. **d**. Jumping probability. **e**. Histogram of pup contact latency. **f**. Violin plots showing the probability distribution of the passive status (status 0) and active status (status 1 and 2). The white dots, boxes, and whiskers represent the median, interquartile range (IQR), and the range extending from 1.5×IQR below the lower quartile to 1.5×IQR above the upper quartile, with whiskers clipped to the data range, respectively. Each data point represents one trial. **g**. Schematic of a GLM model. **h**. Effects of pup and height on behavioral statuses. Color and sizes of circles represent effect weights (GLM coefficients) and significance levels (p value). **i**. Schematic of the risk-taking familiar object contact behavior. **j**. Latency to jump in the risk-taking familiar object contact behaviors. **k**. Histogram of object contact latency. For **c, d**, and **j**, data are mean ± s.e.m. n=7 mice (**c, d**) and n=6 mice (**j, k**). *p<0.05; ****p<0.0001, GLM (**c, d, h** and **j**).

Each day on post-natal day (PND) 3 to 5 at the test stage, eight trial conditions (5, 10, 15, 20 cm height levels × pup presence/absence) were presented to mice in a randomized order in each session. Based on the body part positions labeled by DeepLabCut^20^, we categorized their behaviors into four statuses (Fig. 1a). Status 0 was characterized as inside the platform and mostly associated with immobility. Status 1 was characterized as risk assessment with the head out of the boundary of the platform, and the mouse was likely to be evaluating the height and planning a jump. Status 2 was characterized as jumping attempts, including reaching its forelimbs out and/or hanging its body along the wall to reach the ground. The motivation to jump increased from statuses 0, 1 to 2. Status 3 was characterized as on the ground, including various events such as landing, pup contact, pup retrieval, and nesting. In a typical trial, a mouse on the elevated platform exhibited multiple transitions between statuses 0-2 before jumping down to the ground (Fig. 1b, Supplementary Video 1 and 2).

To evaluate how fear of heights and presence of pups influenced behaviors, we first quantified the latency to jump and the probability of jumping. Greater heights dramatically increased the latency and decreased the probability, while the presence of pups significantly decreased the latency and increased the probability (Fig. 1c, d). After jumping onto the ground, mice contacted pups within 5 s in more than 70% of the pup+ trials, and overall in a short latency (7.75±0.85 s) and with a high probability (95% of the jumping trials) (Fig. 1e, Extended Data Fig. 1a-b). In 66% of the jumping trials, mice retrieved pups within 60 s (time limit set in the experiment) after landing, and the average retrieval latency was 26.03±1.51 s (Extended Data Fig. 1c-b). Given that performances of maternal behaviors in virgins are susceptible to environmental stress^2,21^, the observed pup contact and retrieval latencies were relatively short, suggesting their high motivation towards pups. We also observed qualitatively similar results in dams (Extended Data Fig. 1e-b), with slightly shorter latencies of jumping, pup contact and retrieval, consistent with previous reports that maternal motivation is stronger in dams than experienced virgins^1,8^.

To further elucidate the effects of height and pup on motivational behaviors before landing, we quantified distributions of behaviors in the passive (low motivation, status 0) and active statuses (high motivation, statuses 1 and 2) (Fig. 1f). Using a generalized linear model (GLM) to fit behavioral statuses, we found that pup presence significantly increased the probability of the active status and decreased the probability of the passive status, while the effect of height was the opposite (Fig. 1g, h). Together, these results indicate that while fear of heights reduces the motivation to jump, the presence of pups and maternal motivation dramatically facilitates overcoming such fear.

To ensure that their motivation was specific to pups, but not arbitrary familiar objects, we conducted a control experiment by replacing pups with a familiar object (a centrifuge tube cap that had been placed in the home cage for 1-3 days prior to experiments) on the ground (Fig. 1i and Extended Data Fig. 1l). Despite the effect of height on the latency to jump, there was no difference between the object+ versus the object-trials (Fig. 1j). Furthermore, the object contact latency was longer than that for pups (27.24±1.45s; Fig. 1e, k), indicating that pups, but not neutral and familiar objects, are able to act as a motive to overcome fear.

### mPFC-PAG inhibition suppresses motivation

To understand neural mechanisms that govern risk-taking maternal behaviors, we focused on the mPFC-PAG circuit. Previous studies have shown that the mPFC is a cortical region that mediates a variety of social behaviors, such as social competition^22–24^, social interaction^25^, and social transfer learning^26^. The mPFC sends projections to the PAG^26–30^, a midbrain hub that coordinates numerous innate behaviors, including fear and maternal behaviors^1,31–33^. Therefore, we hypothesized that the mPFC-PAG circuit might act as a high-level control over innate behaviors in complex social contexts.

To determine the causal role of the mPFC – PAG circuit in risk-taking maternal behaviors, we expressed ArchT in the mPFC (see methods) and delivered light to PAG to inhibit axon terminals of the mPFC in the PAG. As a control, we expressed mCherry in the mPFC. For a single mouse, the light+ and light-trials were presented randomly (Fig. 2a). Statistical analysis of the jumping latency (Fig. 2b-b; Extended Data Fig. 2a) and the behavior status distribution (Fig. 2f; Extended Data Fig. 2b-b) both indicated that inactivation of the mPFC-PAG circuit suppressed the effect of pups on the behavior, but without influencing the effect of height. It is likely that other neural circuits, such as the basolateral amygdala (BLA), superior colliculus (SC), and ventral lateral geniculate nucleus (LGN)^18,19^ also mediate the fear of height. Therefore, suppression of the mPFC – PAG circuit alone was insufficient to alter the effect of height. However, this cannot exclude the possibility that mPFC neurons may encode height information (see the next section). Taken together, these results support that the mPFC – PAG circuit is specifically required for the effect of pups on overcoming fear of height during risk-taking maternal behaviors.

**Figure 2.**
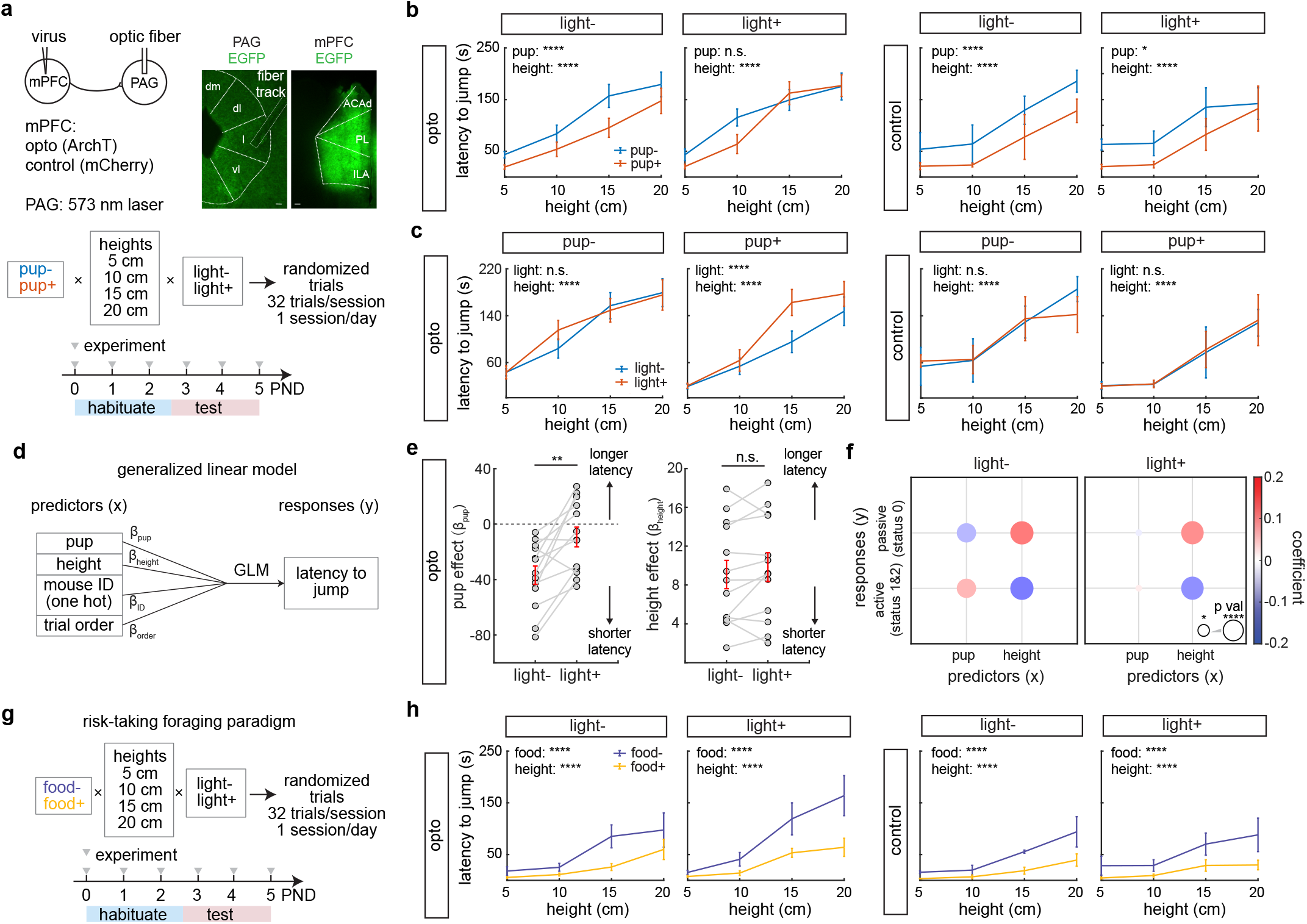
Optogenetic inhibition of the mPFC – PAG circuit suppresses pup-oriented risk-taking behaviors. **a**. Experiment design and representative images of fiber implant and mPFC axon terminals in the PAG and the injection site at the mPFC. Scale bars, 100 µm. dm, dorsomedial; dl, dorsolateral; l, lateral; vl, ventrolateral; ACAd, anterior cingulate area, dorsal part; PL, prelimbic area; ILA, infralimbic area. **b**. Latency to jump with and without light stimulation in the opto and control groups. **c**. Latency to jump in the presence or absence of pups in the opto and control groups. **d**. Schematic of a GLM model predicting the latency to jump. **e**. The pup and height effects measured as predictor coefficients (β) in the GLM model in **d**. Light stimulation diminished the pup effect on the jumping latency, while the height effect remained unchanged. Each pair of gray dots represents data from each mouse. Red lines represent mean ± s.e.m. **p<0.01, rank-sum test. **f**. Effects of pup and height on behavioral statuses in light+ and light-trials. Colors and sizes of circles represent effect weights (GLM coefficients) and significance levels (p value). **g**. The optogenetics paradigm for the risk-taking foraging behaviors. **h** Latency to jump in the risk-taking foraging behaviors with and without light stimulation in the opto and control groups. For **b, c** and **h**, data are mean ± s.e.m. n=13 mice (opto group-pup), n=3 mice (control group-pup) (**b, c, e and f**), n=5 mice (opto group-food) and n=3 mice (control group-food) (**h**). *p<0.05, ****p<0.0001, GLM (**b, c, e, f and h**).

To further test that the mPFC – PAG circuit specifically mediates pup-motivated risk-taking behaviors, we conducted a risk-taking foraging test, in which a food-restricted mouse needed to jump off the elevated platform to get food pellets (Fig. 2g; Extended Data Fig. 2e). In food+ trials, the jumping latency was significantly shorter than food-trials, suggesting that foraging motivation was also able to overcome fear (Fig. 2h). However, when we optogenetically inactivated the mPFC-PAG circuit, the effect of food was unaffected (Fig. 2h; Extended Data Fig. 2f), indicating that the mPFC – PAG circuit is essential to overcome fear during risk-taking behaviors driven by maternal, but not foraging motivation.

### mPFC-PAG encodes motivation

To investigate how motivation is represented in the mPFC-PAG circuit, we next expressed jGCaMP7f in PAG-projecting mPFC neurons, and implanted GRIN lenses in the mPFC for miniscope recording (Fig. 3a). Behavioral performances in the implanted animals were qualitatively similar with the previous results: they showed significant height and pup effects. However, they exhibited longer latencies and lower probabilities of jumping, pup contact and retrieval (Extended Data Fig. 3a-b), probably due to the weights of the setup on their heads. In total, 641 neurons were recorded and analyzed, from 20 sessions in 7 mice. Because behaviors were highly dynamic, we inferred spikes based on calcium signals in order to increase the temporal resolution, and used spike data for all the following analyses (Fig. 3b). Single neurons exhibited diverse activity patterns, responding to a variety of events or behaviors such as wall down (WD, trial initiation), risk assessment (RA), jumping attempt (JA), landing (LA), and pup retrieval (RT) (Fig. 3c; Extended Data Fig. 4a). Typically, neurons that responded to one of the events above were also activated by other events (Fig. 3d), and a significantly higher fraction of cells were activated by multiple behavioral events than chance (see methods; Fig. 3e; Extended Data Fig. 4b). To further compare population activity patterns across various events, we measured correlation coefficients of the population activity vectors between events, and found that activity patterns during IP (inside platform) were distinct from motivational behaviors, such as RA, JA, LA and RT (Fig. 3f). Moreover, overall positively correlated and similar activity patterns were observed between various motivational behaviors (Fig. 3f-b; Extended Data Fig. 4c), suggesting that the mPFC-PAG neurons encode general risk-taking maternal motivation at various behavioral stages.

**Figure 3.**
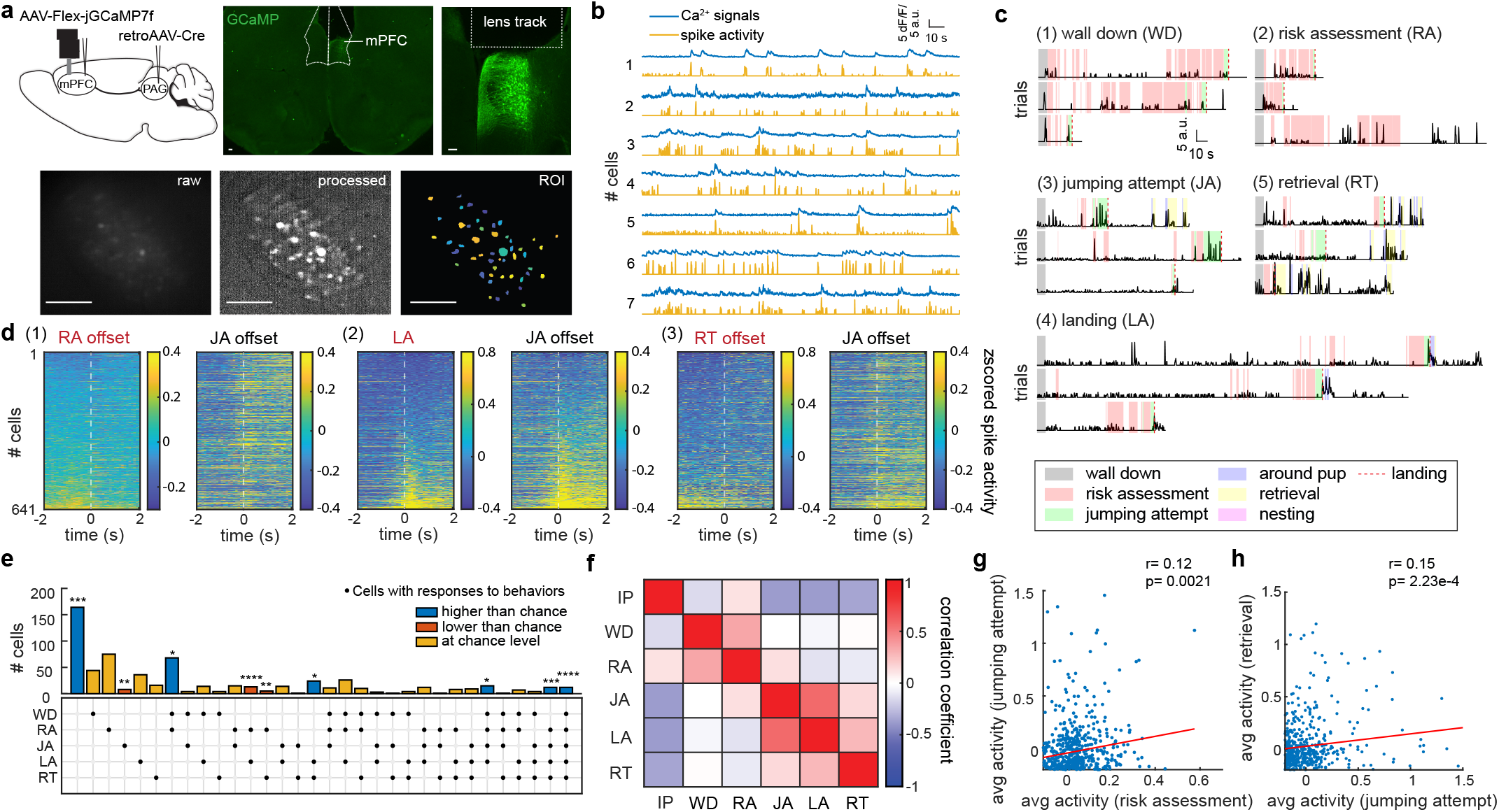
The PAG-projecting mPFC neurons encode general risk-taking maternal motivation at various behavioral stages. **a**. Experiment design (upper left), virus expression and lens track at the mPFC (upper right), and the raw and processed miniscope images in the same field of view (bottom). Scale bars,100 µm. **b**. Calcium signals (dF/F, blue) and the corresponding inferred spike activity (yellow) of example simultaneously recorded cells. **c**. Spike activity of five example neurons each consistently responding to a behavioral event. **d**. Heatmaps of z-scored spike activity across all recorded neurons aligned with three pairs of events. Neurons were sorted by the average signal amplitude across the entire duration of a behavior labeled in red. **e**. The number of neurons significantly activated during different behaviors, compared to a null model in which neuronal response to each behavior is independent. Bottom, each column represents a neuronal activation pattern, and each row represents a behavioral event. Dots indicate that neurons with a given activation pattern are activated in certain behaviors. See methods for statistical analysis. *, **, *** and **** denote Bonferroni corrected p values < 0.05, 0.01, 0.001 and 0.0001 respectively. **f**. The population activity correlation matrix between behaviors. **g-h**. Scatter plots showing activity during risk assessment versus jumping attempt (**g**), or during jumping attempt versus retrieval (**h**). Each dot represents an individual neuron. The x or y axis represents average activity during a certain behavior. Red lines represent the fitted lines. r, Pearson correlation coefficients. p, p values. IP, WD, RA, JA, LA, and RT denote inside platform, wall down, risk assessment, jumping attempt, landing and retrieval events respectively (**c**-**f**). n=641 cells from 20 sessions in 7 mice (**d-h**).

To further elucidate the representation of motivation in the mPFC-PAG circuit, we focused on the activity during risk assessment and/or jumping attempts. In theory, cells activated during these events might represent motivation, fear, or vestibular signals (e.g. hanging on the wall). Our inactivation and imaging results strongly suggested involvement of this circuit in motivation because: (1) inhibition of the mPFC-PAG circuit results in lower motivation in risk-taking maternal behaviors as in Fig. 2; and (2) the mPFC-PAG neurons are activated during multiple motivational behaviors such as risk assessment, jumping attempts and pup retrieval as in Fig. 3d-b. To further validate the motivation model, we compared activity across conditions, and expected distinguishable results across the three models mentioned above (motivation model, fear model, and vestibular model; Fig. 4a). In the motivation model, the activity will be higher with pup presence and lower heights, and also stronger in a successful jumping attempt followed by landing (pre-jump) than a failed jumping attempt followed by staying inside the platform or risk assessment (pre-retreat). In the fear model, the activity will be higher with greater heights, stronger in a pre-retreat attempt, and not affected by the presence of pups. In the vestibular signal model, the jumping attempt activity will be higher with greater heights, stronger in a pre-jump threat, and not affected by the presence of pups.

**Figure 4.**
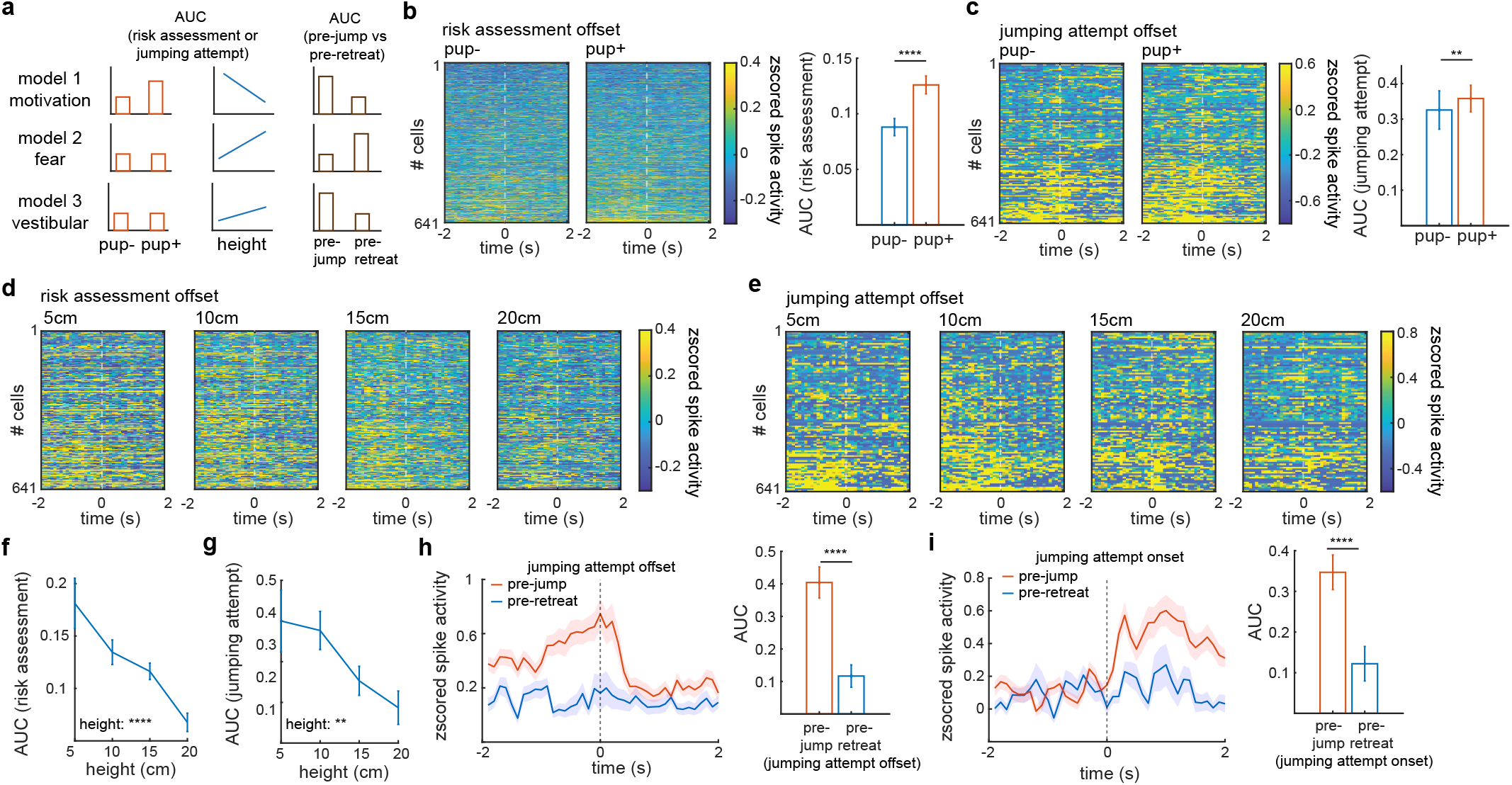
The PAG-projecting mPFC neurons encode motivation, but not fear or vestibular information. **a**. Distinct predictions on neural activity during risk assessment or jumping attempts by motivation, fear and vestibular sensation models. **b**. Left, heatmaps of cross-validated z-scored spike activity aligned with the offset of risk assessment. Neurons are sorted by their activity during the entire event of risk assessment. Right, a bar graph comparing the activity of the risk assessment-activated neurons during the entire risk assessment event between pup-versus pup+ trials. ****p<0.0001, signed rank test. **c**. Same as **b**, but for jumping attempts. **p<0.01, signed rank test. **d-e**. Heatmaps of cross-validated z-scored spike activity aligned with the risk assessment offset (**d**) and the jumping attempt offset (**e**) at different heights. Neurons are sorted by their activity during the entire events. **f**. Average activity of the risk assessment-activated neurons during the entire risk assessment event. at different heights. ****p<0.0001, GLM. **g**. Average activity of the jumping attempt-activated neurons during the entire event of jumping attempts at different heights. **p<0.01, GLM. **h**. Left, average z-scored spike activity of the jumping attempt-activated neurons aligned with the jumping attempt offset in pre-jump and pre-retreat trials. Right, a bar graph comparing average activity of the jumping attempt-activated neurons in the 2 s window before the offset for the pre-jump and pre-retreat trials. ****p<0.0001, signed rank test. **i**. Same as h, but for the jumping attempt onset. Data are mean ± s.e.m. (**b, c**, and **f-i**). See Supplementary Information for the number of cells for each panel. Cells were from 20 sessions in 7 mice (see methods).

To test the three proposed models, we compared neural activity during risk assessment or jumping attempts across different trial conditions and found that neuronal responses significantly increased in the presence of pups (Fig. 4b, c; Extended Data Fig. 4d, e) and decreased with greater heights (Fig. 4d-b; Extended Data Fig. 4f, g). Furthermore, the pre-jump responses were dramatically larger than the pre-retreat responses during jumping attempts (Fig. 4h, i). Interestingly, we also noticed a height-dependent decrease in pup retrieval activity (Extended Data Fig. 4h, i), which might reflect reduced maternal motivation after experiencing more stresses from height (Extended Data Fig. 3d-b). These data, together with the optogenetic results, support a model that the PAG-projecting mPFC neurons encode motivation in the risk-taking maternal behaviors, and drive diverse pup-directed behaviors including risk assessment, jumping attempts, and pup retrieval.

### Representational geometry allows for cue integration

Having shown that the motivation to jump is modulated by fear of height and presence of pups (Fig. 4b-b), we next asked how the pup and height cues are integrated in the mPFC-PAG circuits to drive behaviors (Fig. 5a). We hypothesized that representational geometry of neural activity space underlies cue integration. Therefore, we focused on two questions: how do pup and height cues modulate neural activity? How does neural activity drive risk-taking behaviors? One intuitive approach is to use a GLM to predict neural activity with both sensory (pup and height cues) and motor (behavior status) variables. However, behaviors are modulated by pup and height cues and hence their correlations (Fig. 1h) might obscure the interpretation of fitting parameters. Moreover, unlike most tasks with delayed responses^4,5^, it is difficult to temporally dissociate sensory and motor variables in naturalistic behavioral paradigms due to their overlaps in time. Therefore, we proposed to address these questions by performing GLM using two generative models.

**Figure 5.**
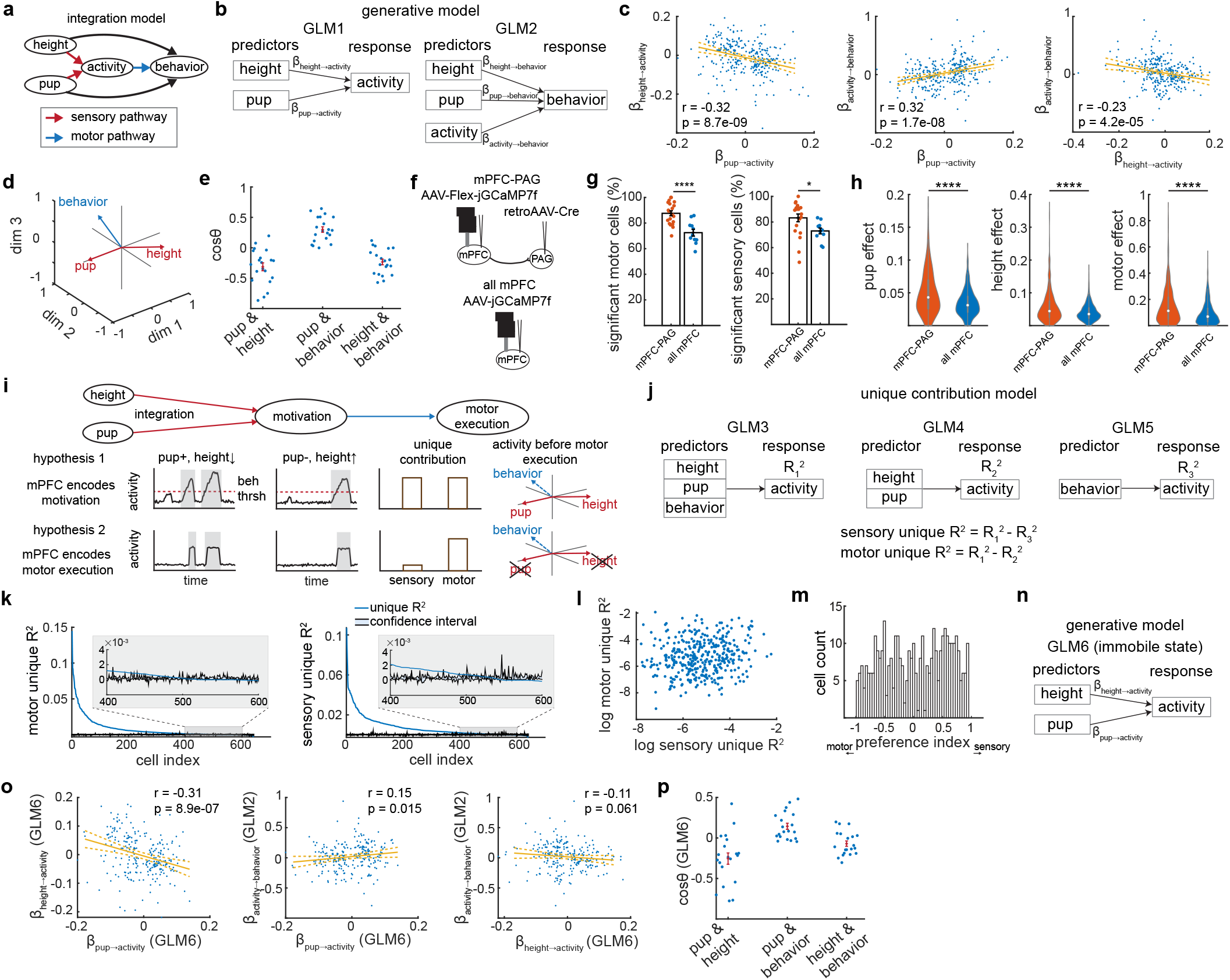
Representational geometry allows for integration of pup and height cues to drive pup-oriented risk-taking behaviors. **a**. Schematic of the integration model in which pup and height cues are integrated in the mPFC-PAG circuit to drive risk-taking maternal behaviors. **b**. Two generative models (GLM1 and GLM2), in which coefficients of the predictors (*β*) were determined. **c**. Scatter plots of *β*_*height*→*activity*_ versus *β*_*pup*→*activity*_ (left), *β*_*activity*→*behavior*_ versus *β*_*pup*→*activity*_ (middle), and *β*_*activity*→*behavior*_ versus *β*_*height*→*activity*_ (right). Each dot represents data from an individual neuron. Yellow lines represent fitted curves with 95% confidence intervals. r, Pearson correlation coefficients. p, p values. **d**. The coding directions of pup, height and behavior, measured as vectors of coefficients from GLM1 and GLM2. The vectors embedded in three-dimensional space by QR matrix decomposition. **e**. Cosine values of angles between coding directions. **f**. Schematic of miniscope recording in the PAG-projecting mPFC neurons versus the entire mPFC. **g**. Percentage of motor and sensory neurons with significant coefficients (non-zero coefficient) in *β* _*activity*→*behavior*_ (left), and *β*_*pup*→*activity*_ or *β*_*height*→*activity*_ (right) respectively. *p<0.05, ****p<0.0001, two-sample *t*-test. **h**. Violin plots showing distributions of absolute values of pup (*β*_*pup*→*activity*_), height (*β*_*height*→*activity*_) and motor (*β* _*activity* → *behavior*_) effects. The white dots, boxes and whiskers represent the median, interquartile range (IQR), and the range extending from 1.5×IQR below the lower quartile to 1.5× IQR above the upper quartile, with whiskers clipped to the data range respectively. ****p<0.0001, two-sample t-test. **i**. A model depicting the process in which pup and height cues are integrated to drive motivation. In hypothesis 1, the mPFC-PAG circuit encodes the motivation (internal state). In hypothesis 2, the mPFC-PAG circuit directly controls motor execution. The diagram also illustrates distinct predictions of the two hypotheses. **j**. Three GLM models (GLM3, GLM4 and GLM5) for measuring sensory and motor unique R^2^. **k**. Motor (left) and sensory (right) unique R^2^ of recorded neurons. **l**. Scatter plot of motor and sensory unique R^2^ of individual neurons with significant motor and sensory unique R^2^ in **k. m**. Distribution of the preference index measured as (sensory unique R^2^ – motor unique R2)/ (sensory unique R^2^ + motor unique R^2^). An index of 1 or -1 indicates absence of motor or sensory unique R^2^ respectively. **n**. A generative model (GLM6) to predict activity during immobile states (status 0) on the elevated platform. **o**. Scatter plots of *β* coefficients from GLM6 and GLM2. Each dot represents data from an individual neuron. Yellow lines represent fitted curves with 95% confidence intervals. r, Pearson correlation coefficients; p, p values. **p**. Cosine values of angles between coding directions in GLM6 and GLM2. Data are mean ± s.e.m. (**e, g**, and **p**) See Supplementary Information for the number of cells for each panel. PAG-projecting mPFC neurons and pan mPFC neurons were from 20 sessions in 7 mice and from 11 sessions in 3 mice respectively (see methods).

In GLM1, we predicted neural activity using pup and height variables to answer how pup and height cues modulate neural activity (red arrows in Fig. 5a; Fig. 5b). In GLM2, we predicted behavior statuses (passive status 0 with low motivation versus active status 1/2 with high motivation) using both pup and height cues and neural activity (Fig. 5b). The coefficients of neural activity in GLM2 represent how activity drives behaviors (blue arrow in Fig. 5a). The coefficients of pup and height cues represent their behavioral effects mediated by other pathways or neurons not recorded in the current experiment (black arrows in Fig. 5a; Fig. 5b; note that their behavioral effects mediated by the mPFC-PAG circuits are reflected in the coefficients of neural activity). Thus, *β*_*height*→*activity*_ and *β*_*pup*→*activity*_ in GLM1 characterize the cue-modulated directions, and *β* _*activity*→*behavior*_ in GLM2 characterizes the motivation-driving direction in the neural space (Fig. 5b). We found that at the population level,, *β*_*height*→ *activity*_ and, *β*_*pup*→*activity*_ were negatively correlated (Fig. 5c-b, Supplementary Video 3), implying antagonistic effects of height and pup cues. Moreover, *β*_*activity*→*behavior*_ is positively correlated with *β* _*pup*→ *activity*_, but negatively correlated with *β*_*height*→ *activity*_ (Fig. 5c-b). These results revealed the representational geometry of the activity space, supporting a model that the presence of pups is able to drive the population activity vector along the direction encoding higher motivation, while increased heights do the opposite. This allows for integration of pup and height information to drive risk-taking maternal behaviors (Fig. 1c). We also recorded and analyzed non-specific mPFC neurons, and found that the fraction of sensory (modulated by pup and height cues) or motor (modulating motivation) cells and their average modulating effects were smaller compared to PAG-projecting mPFC neurons (Fig. 5f-b), further suggesting the specificity of the mPFC-PAG pathway in this task.

We next examined whether the mPFC-PAG circuit directly controls motor execution, or represents an internal state that integrates environmental cues and gives rise to motivation (Fig. 5i). While in both models we expected to see a positive correlation between *β* _*activity*→ *behavior*_ and *β*_*pup*→*activity*_, and a negative correlation between *β* _*activity*→ *behavior*_ and *β*_*height*→ *activity*_, previous results in which the activity during risk assessment or jumping attempts are higher with pup presence and lower heights in Fig. 4 strongly suggested the internal state hypothesis. Here we performed two additional analyses to test the hypothesis. In the first analysis, we measured the unique contributions of the sensory or motor variable to neural activity^34^ (Fig. 5j). We predicted that if the mPFC-PAG circuit encodes motor execution, its activity will be mostly explained by motor variables, with minimal unique contributions from sensory variables. However, our results showed that both sensory and motor variables made significant unique contributions to predicting most of the neural activity (Fig. 5k-b; Extended Data Fig. 5). In the second analysis, we focused on the activity before motor execution, i.e. when the animals were immobile on the platform (Fig. 5n). We reasoned that if the mPFC-PAG circuit represents the internal state, the motivation should emerge even before the animal takes actions. Indeed, we found that neural activity was already influenced by pup and height cues in opposite directions in the immobile state (Fig. 5o, p), and the direction modulated by pup in the immobile state was positively aligned with the motivation-driving direction in GLM2 (Fig. 5o, p). Hence, our findings suggested that the representational geometry of the mPFC-PAG activity mediates the internal process in which environmental cues are integrated to generate motivational behaviors.

### Switching dynamics drives motivation

In the mPFC, integration of pup and height cues gives rise to risk-taking maternal motivation. However, the dynamics of such motivation remains unclear. Previous results have reported that a continuous ramping model is able to explain dynamics of motivation or decision variables in certain behaviors. In this model, the activity continues to ramp in a certain dimension, and elicits behaviors/decisions after reaching a threshold (Fig. 6a). This type of dynamics is typically observed during structured tasks, such as perceptual decision making, in which well-trained animals need to make decisions in short time windows after cue presentation^4,5^, or during natural behaviors such as escaping^32^, aggression and mating^35–37^, in which behaviors are elicited by strong motivation arising from sudden and salient cues (e.g. presentation of looming stimuli, or introduction of conspecifics) in situations with little or no conflict. However, animals could display a variety of spontaneous behaviors in ethological contexts, characterized by stochastic and less-predictable transitions between multiple possible behaviors in the behavioral repertoire, such as exploration, grooming, or sleep^38–41^. Here in our risk-taking maternal behavior test, animals also exhibited spontaneous behaviors in this semi-naturalistic and conflicting situation: in most trial conditions, animals on the elevated platform exhibited multiple stochastic transitions between immobility, risk assessment, and jumping attempts, and the moment of jumping was not precisely locked to any external cues (Fig. 1b). In such contexts, external cues are more likely to influence, but not immediately evoke behaviors. To explain neural dynamics that gave rise to such behaviors, we proposed a switching ramping model, in which the activity switches between a ramping state and a non-ramping state, resulting in fluctuating motivation and hence stochastic transitions between spontaneous behaviors.

**Figure 6.**
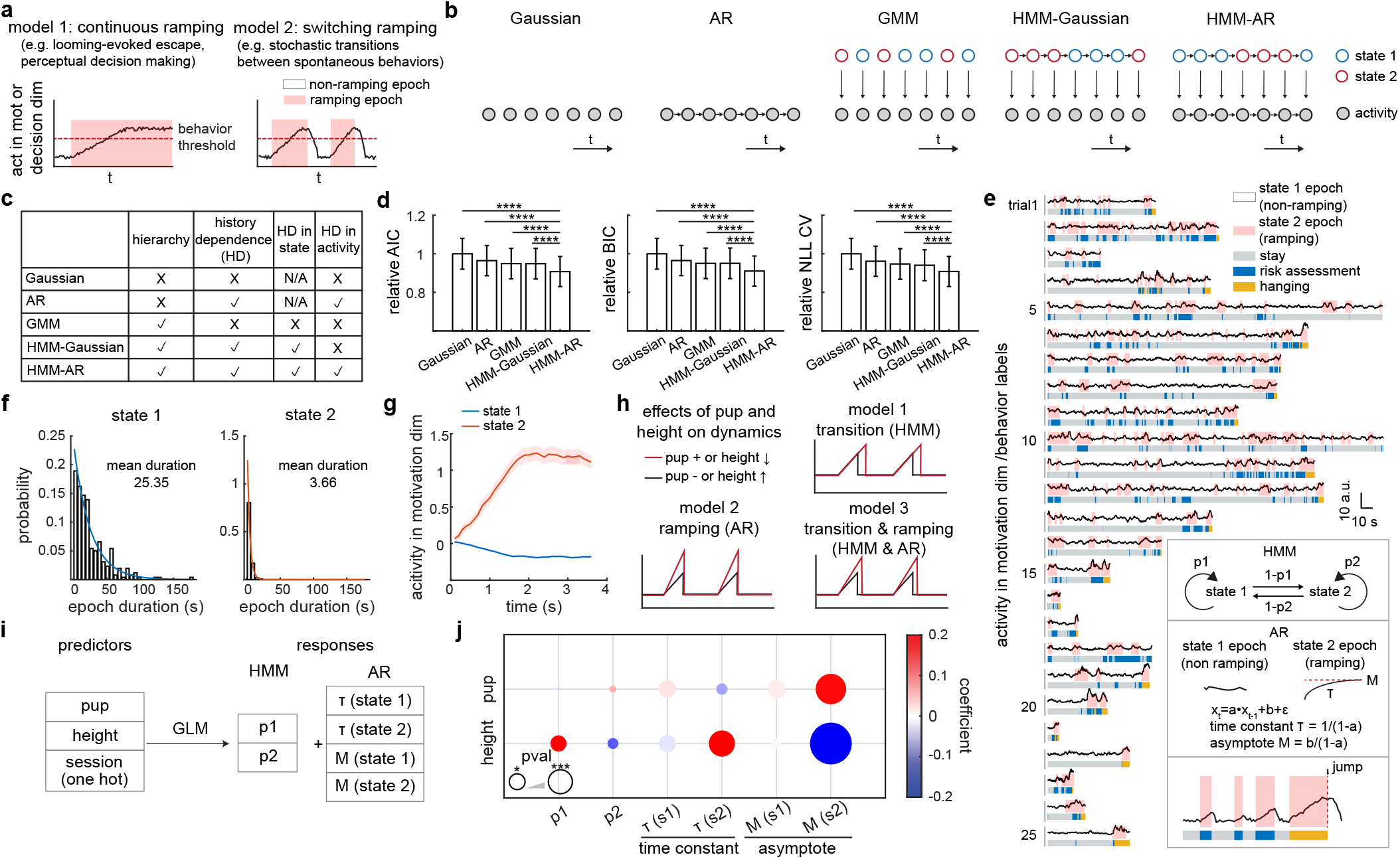
Switching dynamics revealed by the HMM-AR model modulates spontaneous pup-oriented risk-taking behaviors. **a**. Two models of neural dynamics underlying how motivation elicits behaviors. **b-c**. Five proposed generative models (**b**) and their characteristics (**c**). **d**. Model comparisons using relative AIC (left), BIC (middle) and NLL CV (right) among five models. The HMM-AR model with the smallest values in all three parameters showed the best performance. ****p<0.0001, signed rank test. **e**. Example trials showing state transitions predicted by the HMM-AR model (white and pink shadows), in together with neural activity in the motivation dimension (upper traces) and behaviors (bottom plots). Inset shows key parameters of the HMM-AR model and the predicted results on the activity. **f**. Distributions of epoch durations for state 1 (left) and state 2 (right) from an example session. **g**. Mean activity in the motivation-encoding dimension during state 1 and 2 epochs from an example session. **h**. Three models in which pup and height cues influence motivational behaviors. **i-j**. A GLM model (**i**) and the effects of predictors (pup and height) on the HMM-AR model parameters (**j**). Colors and sizes of circles represent effect weights (GLM coefficients) and significance levels (p value). *p<0.05, ***p<0.001, GLM. Data are mean ± s.e.m. (**d, g**). n=641 neurons from 20 sessions in 7 mice.

To systematically quantify neural dynamics, we proposed a series of generative models, and evaluated their performance in predicting neural activity dynamics in the motivation-encoding dimension defined by *β* _*activity*→ *behavior*_ in GLM2 in Fig. 5b. These models include a Gaussian model, an autoregressive model (AR), a Gaussian mixture model (GMM), a hidden Markov – Gaussian model (HMM-Gaussian), and a hidden Markov – autoregressive model (HMM-AR) (Fig. 6b)^42^. These models differ in whether they are hierarchical or whether they are history-dependent (HD) (Fig. 6c). In hierarchical models such as GMM, HMM-Gaussian and HMM-AR, distinct states in the higher hierarchy result in distinct activity dynamics in the lower hierarchy. In state-HD models such as HMM-Gaussian and HMM-AR, the Markovian property of state transitions might result in block-like organization of states. In activity-HD models such as AR and HMM-AR, the autoregressive property of activity might give rise to ramping dynamics. Therefore, the Gaussian model represents the simplest model without hierarchical or temporal structures, while the HMM-AR model incorporates the most elaborate structures, potentially allowing for the switching ramping dynamics.

Model comparison results showed that the HMM-AR model best predicted the observed neural dynamics in the motivation-encoding dimension, as measured by Akaike information criterion (AIC), Bayesian information criterion (BIC), or cross-validated negative log-likelihood (NLL CV) (Fig. 6d). Moreover, as predicted by the HMM-AR model, the dynamics could be explained by switches between a ramping state (state 2) and a non-ramping state (state 1) (Fig. 6e). In general, the non-ramping state (state 1) resulted in lower activity and lower motivation (more status 0), and showed a longer duration. Conversely, the ramping state (state 2) resulted in ramping activity and higher motivation (more status 1 or 2), and showed a shorter duration (Fig. 6e-b). It is worth noting that the ramping activity was unlikely a result of the slow dynamics of calcium signals, because: (1) the spike inference algorithm that we used for data processing has already been optimized to preserve auto-correlation of real spikes^43^; and (2) auto-correlation in the motivation-encoding dimension was much higher than that in a random dimension, suggesting that the high auto-correlation and hence ramping activity in the motivation-encoding dimension could not simply explained by imperfect deconvolution of calcium signals (Extended Data Fig. 6a). Altogether, our results support that the switching ramping feature of neural activity underlies the spontaneity of neural dynamics and behaviors.

As pup and height cues were able to influence behaviors, we next asked how the switching ramping dynamics is modulated by environmental cues: they might influence state transition probabilities (model 1), ramping parameters (model 2), or both (model 3) (Fig. 6h). To answer this question, we used a GLM model to predict condition-specific HMM parameters (transition probabilities p1 and p2 in the HMM model) and AR parameters (time constant *τ* and asymptote M in the AR model) with predicting variables including heights and presence of pups (Fig. 6i). We found a trend of shorter durations of state 1 epochs (non-ramping state) and longer durations of state 2 epochs (ramping state) with pup presence or at lower heights (Fig. 6j; Extended Data Fig. 6b, c). Additionally, the ramping parameters, especially for state 2, including the time constant and asymptote were significantly and strongly modulated by environmental cues. Increased heights resulted in smaller ramping asymptotes, slower time constants and hence lower ramping amplitudes in the motivation-encoding dimension (Fig. 6j; Extended Data Fig. 6e), in line with lower motivation and longer jumping latencies observed in behaviors. In contrast, the presence of pups resulted in higher ramping amplitudes, giving rise to higher motivation and shorter jumping latencies (Fig. 6j; Extended Data Fig. 6d). Taken together, pup and height cues modulate the switching ramping dynamics to influence, but not immediately evoke behaviors, prominently via modulating dynamics during the ramping state.

## Discussion

Our results show that maternal motivation is able to overcome innate fear and drive risk-taking behaviors, specifically mediated by the mPFC-PAG circuit (Extended Data Fig. 7a). Representational geometry in the circuit allows for integrating conflicting cues about pups and heights to encode risk-taking maternal motivation. Moreover, animals display spontaneous and stochastic transitions between different behavioral statuses in this semi-naturalistic paradigm with conflict, arising from the switching ramping dynamics in the circuit, which is further modulated by external cues to influence motivation and behaviors. Altogether, our findings highlight the essential role and computational mechanism of the prefrontal-brainstem pathway in resolving the conflict between maternal motivation and innate fear, which might advance our understanding of animal sociality and altruism.

Previous studies have shown that innate defensive behaviors, including fear of heights, are mostly mediated by subcortical regions such as the amygdala, hypothalamus, and PAG^18,19,31,32,44,45^. Our results show that with inhibition of the mPFC-PAG circuit, fear of heights still existed, but animals were no longer driven by maternal motivation to overcome such fear. In relation to this, recent studies also reported critical roles of the mPFC-subcortical circuits in modulating innate defensive or foraging behaviors in complex social contexts, such as observational learning, and food competition^22,23,26^. Hence, we speculate that while evolutionarily conservative subcortical regions might be sufficient for most innate behaviors that are essential for survival and reproduction, the descending projections from the neocortex to the subcortical regions provide social control over innate behaviors, allowing for additional flexibility of adaptive behaviors and emergence of animal sociality in complex social contexts.

In the risk-taking maternal behavior paradigm, we observed spontaneous behavior patterns characterized by stochastic transitions between statuses, the timings of which could not be precisely predicted by external cues. Our analyses show that the switching ramping model best explains the mPFC dynamics, allowing for probabilistic transitions between a non-ramping state and a ramping-state, resulting in the observed behavioral patterns. However, neural mechanisms that underlie such state switching remain unclear. Future work will be needed to reveal whether such state switching is global across the entire cortex, or local to individual areas. The roles of neuromodulators, which are crucial for the ON-OFF switching in sensory areas^46,47^, should also be inspected in this paradigm. Moreover, we propose to understand neural dynamics in other naturalistic environments, and we speculate that the switching ramping dynamics might represent a general model that gives rise to spontaneous behaviors.

## Methods

### Animals

All procedures were approved by the Institutional Animal Care and Use committee at the Institute of Biophysics, Chinese Academy of Sciences and performed in accordance with institute’s guidelines for the care and use of laboratory animals. Wild-type C57BL/6J female mice aged 8-12 weeks were used as test subjects in this study. Pups aged post-natal day 0-5 were from wild-type C57BL/6J mouse pairs. Mice were housed under a 12 h light/12 h dark cycle with food and water available *ad libitum*. Mice were randomly selected for the test and the control groups.

### Histology

Mice were transcardially perfused with 4% paraformaldehyde (PFA; Biosharp, BL539A). Brains were dissected, post-fixed in 4% PFA overnight at 4 °C and then transferred to phosphate-buffered saline (PBS; Biosharp, BL551A). Brains were sliced at 50 μm on a Leica VT1000 S vibratome. For immunostaining, free-floating brain sections were washed with PBS 3 times, followed by blocking in 5% normal goat serum (NGS; Beyotime, C0265) and 0.1% Triton X-100 (Beyotime, ST97) for 1 h at room temperature (RT). Sections were incubated with primary antibody (rabbit anti-green fluorescent protein, 1:1000; Invitrogen, A11122), 0.5% NGS and 0.1% Triton X-100 overnight at 4 °C. The next day sections were incubated with secondary antibody (Alexa Fluor 488 goat anti-rabbit IgG, 1:1000; Invitrogen, A11008), 0.5% NGS and 0.1% Triton X-100 at RT. Sections were washed with PBS 3 times and then mounted on coverslips with DAPI Fluoromount-G (SounthernBiotech, 0100-20).

### Imaging and image analysis

Brain sections were imaged under a Leica MZ10 F modular stereo microscope and a Leica DM2500 LED optical microscope. Images were analyzed with Leica LAS X software.

### Viruses

The following commercially available viruses were used in this study: AAV-CAG-FLEX-ArchT-EGFP-WPRE (5.1×10^12^ GC/mL, OBiO, AG28307), AAV-CAG-ArchT-GFP (6.1 ×10^12^ GC/mL, OBiO, AG29777), AAV-hSyn-DIO-mCherry-WPRE (5.5×10^12^ GC/mL, OBiO, H4828), AAV-CMV-NLS-Cre-WPRE (5.5×10^12^ GC/mL, OBiO, CN0998), retroAAV-hSyn-Cre-WPRE (1.8×10^12^ GC/mL, OBiO, CN867), AAV-hSyn-FLEX-jGCaMP7f-WPRE (1×10^12^ GC/mL, OBiO, H11257), AAV-hSyn-jGCaMP7f-WPRE (2.1×10^12^ GC/mL, OBiO, H11265). All viruses were aliquoted on the day of delivery and stored at -80°C until use.

### Stereotaxic surgery

Mice were intraperitoneally injected with a mixture of tribromoethanol, meloxicam and Baytril prior to surgeries. Animals were placed at the stereotaxic rig and anesthetized with 1-2% isoflurane in oxygen at a flow rate of 1-2 L/min. Viruses were injected into the brain with a nanoinjector (RWD, R480).

For optogenetics experiments, three strategies of ArchT virus injection (nL per hemisphere) were used and made no differences in behavioral performances: (1) 200 nL of AAV-CAG-ArchT-GFP in mPFC; (2) 100 nL of AAV-CAG-FLEX-ArchT-EGFP-WPRE injected in mPFC, and 80 nL of retroAAV-hSyn-Cre-WPRE injected in PAG; (3) 200 nL of a 1:1 mixture of AAV-CMV-NLS-Cre-WPRE and AAV-CAG-FLEX-ArchT-EGFP-WPRE in mPFC. For the control experiment, 200 nL of a 1:1 mixture of AAV-CMV-NLS-Cre-WPRE and AAV-hSyn-DIO-mCherry-WPRE was injected in mPFC. Viruses were bilaterally injected into mPFC (AP: +1.65, ML: +/-0.5, DV: -1.5 from brain surface) and/or PAG (AP: -0.55 from lambda, ML: +/-1.0, DV: -2.45 from brain surface, 18°). Optical fiber cannulas (200 μm in diameter, NA 0.37, 4 mm in length) were bilaterally implanted in PAG (AP: -0.55 from lambda, ML: +/-1.0, DV: -1.8 ∼ -2.4 from brain surface, 18°) and secured to the skull with dental cement (Nissin, 1R).

For miniscope experiments, to label PAG-projecting mPFC neurons, 400 nL of AAV-hSyn-FLEX-jGCaMP7f-WPRE and 150 nL of retroAAV-hSyn-Cre-WPRE were injected in mPFC (AP: +1.65, ML: + 0.5, DV: -1.5 from brain surface) and PAG (AP: -0.55 from lambda, ML: +1.0, DV: -2.0 from brain surface) respectively. To label mPFC neurons non-specifically, 300 nL of AAV-hSyn-jGCaMP7f-WPRE was injected in mPFC (AP: +1.65, ML: + 0.5, DV: -1.4 from brain surface). The scalp skin was sutured to protect the skull. Animals were allowed to recover from surgery for at least two weeks before viruses were fully expressed. To implant the gradient refractive index (GRIN) lens, a circular craniotomy with a diameter of ∼1.5 mm was made around the target site of mPFC (AP: +1.65, ML: +0.5). Dura was carefully removed with fine forceps and a bent needle. Brain tissues (DV: -1.1 from brain surface at the deepest) were vacuumed out with a 26-gauge blunt syringe needle attached to a stereotaxic arm. A GRIN lens (0.433 pitch, 1.0 mm diameter and 3.758 in length; Go!Foton, CLHS100GFT003) was attached to a customized holder connected to vacuum and slowly implanted to the target depth (DV: -1.5). Lens was secured in place firstly by 3M Vetbond (3M, 1469) and subsequently by Metabond dental cement (Metabond, C&B, S380). Next a UCLA miniscope v4.4 (OpenEphys, OEPS-7407) was fitted with a baseplate, attached to a holder and lowered down to a plane 0.2 mm above the surface of the GRIN lens. The baseplate was secured with dental cement. When not in use, a customized dummy miniscope was placed in the baseplate to protect the GRIN lens from dust and for animals to get used to the weight on their heads. Animals were returned to their home cages and recovered for at least three weeks before experiments.

### Behavioral experiments

Behaviors were recorded at 30 frames/s with a Logitech webcam on top of the customized behavior apparatus in a sound-attenuated chamber. The behavioral apparatus consisted of a 52 cm × 52 cm arena (ground) and a 10 cm × 10 cm elevated platform in the middle with a surrounding barrier connected to a motor driver and a controller that controlled its rise and fall. The elevated platform could be set at four different heights (5, 10, 15 and 20 cm) from the ground by adjusting the position of the arena. Behavioral experiments were conducted during animals’ light phase (9:00h – 19:00h).

In risk-taking maternal behavior assays, each female virgin mouse was introduced to a pregnant mouse and co-housed with the dam and her pups through postnatal day (PND) 5. PND 0 – 2 was the habituation period with 24 trials/session and 1 session/day so that test animals could get used to the behavioral apparatus and learn to jump from the elevated platform. PND 3 – 5 was the test period with 32 trials/session and 1 session/day. Data from PND 3 – 5 were pooled together for analysis. On each day, mice were allowed to habituate with 3 – 5 pups on the ground of the behavior apparatus with their nest in a corner for 10 min at the beginning of experiments. Eight trial conditions in combinations of pup (presence or absence) and height (5, 10, 15, 20 cm) variables were randomized. Each condition was replicated three and four times on each day of the habituation period and the test period respectively. At the beginning of each trial, the test animal was placed on the elevated platform with the surrounding barrier raised up so that the view of the context such as presence of pups and distance to the ground was blocked. In pup present trials, pups were scattered away from the nest. The height of the elevated platform was adjusted by moving up or down of the ground platform. When a trial started, the barrier was lowered down and the mouse was able to explore the edges of the elevated platform to observe the surroundings. The entire trial took no more than 5 min. If the mouse jumped off the elevated platform before 5 min, it would have about 1 min on the ground to contact or retrieve pups, build nest or explore etc. Trials in which mice jumped were included for the analysis of the distribution of pup contact or retrieval latency. Mice for optogenetics or miniscope experiments followed the same behavioral paradigm and were habituated for at least 3 days beforehand to get used to the weight on their heads.

In risk-taking familiar object contact assays, a green-color cap of a 15 mL conical tube was placed in the home cage of virgin female mice for 1-3 days before experiments. The cap was returned to the home cage with test animals after experiments throughout test day 1 – 6. Each day had 32 trials/session and 1 session/day. The experiment procedures in the behavior apparatus were the same as the risk-taking maternal behavior assays except that the pups were replaced with the familiar cap.

In risk-taking food consumption assays, virgin female mice were food deprived for 20 h every day before experiments. The experiment procedures in the behavior apparatus were similar to the risk-taking maternal behavior assays except that the pups were replaced with a food pellet and the test animals were allowed to consume food after landing on the ground for about only 15 s. After experiments the test animals had free access to chow food for 30 – 60 min in the home cage.

### Behavioral annotation and tracking

Animal behaviors were annotated offline with BORIS^48^ (Behavioral Observation Research Interactive Software). Wall down was defined as the moment when the barrier around the elevated platform completely came down. Landing was defined as the first frame in which all the limbs of animals landed on the ground. Pup contact was defined as the frame in which test animals first contacted a pup on the ground after landing. Retrieval was defined as the first frame in which test animals picked up a pup.

Animal positions on the elevated platform in behavior (Fig. 1) and optogenetics (Fig. 2) experiments were tracked with DeepLabCut^20^. Based on the position of their labeled body parts (head, the anterior, middle and posterior parts of the body, and tail base), four states of behaviors were categorized to represent their motivation of jumping off the elevated platform: (1) status 0, all labeled body parts were restrained inside the elevated platform; (2) status 1, only the head stretched out of the elevated platform; (3) status 2, all body parts other than the tail base were out of the elevated platform before landing defined by BORIS; (4) status 3, all body parts were out of the elevated platform after landing.

To align behavior labeling with neural data in fine resolution, we manually annotated animal behaviors on the elevated platform with BORIS in the miniscope experiments (Fig. 3-6; Extended Data Fig. 4-6). Risk assessment was defined as frames with test animals’ head out of the elevated platform followed by stretching their head and neck out in the beginning. Jumping attempt was defined as frames with all body parts hanging on the wall prior to landing. Around pup was defined as frames in which animals exhibited pup-oriented behaviors on the ground. Retrieval was defined as frames from picking up a pup to dropping a pup in nest. Nesting was defined as frames in which animals built a nest.

### Optogenetics

The bilateral fiber cannulas were connected to a splitter branching optical fiber and a rotary joint (CNI-Laser). Constant 10 mW light stimulus per hemisphere was delivered by a 594-nm laser (CNI-Laser, FC-589nm) which was controlled by a Pulse Pal (Sanworks, v2) and Bonsai^49^ on a computer. The risk-taking maternal behavior assay with optogenetic manipulation was similar to the one without. However, other than the pup and height variables, light stimulus (with or without light) was the third variable in the randomized trial design. Each combination of the three variables was replicated twice on each day of the test period.

### Behavioral analysis using generalized linear models

Generalized linear models (GLM) were employed to investigate the effects of pup, object or food and platform height on behaviors, such as latencies or probabilities of jumping, pup contact, or pup retrieval (Fig. 1c, d, j; Fig. 2b, c, e, h; Extended Data Fig. 1a-b; Extended Data Fig. 2a, f; Extended Data Fig. 3a, b, d-g), and probabilities of behavioral statuses (Fig. 1h; Fig. 2f; Extended Data Fig. 2d). The model is defined as:

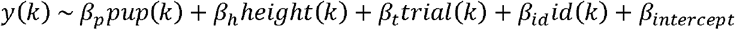

The predictors include: (1) *pup* (*k*), which denotes the presence of pups, which is one-hot encoded. Z-score standardization was applied subsequently to predict probabilities of behavioral statuses. (2) *height* (*k*), which denotes z-score standardized platform height. (3) *trial* (*k*), which denotes the trial number. (4) *id* (*k*), which denotes individual mouse identity, encoded as one-hot variables. The coefficients *β* quantify the effects of these predictors on the response variable. When predicting latencies of jumping, pup contact or pup retrieval, *y* (*k*) denotes the latency in trial *k*, and a normal distribution with an identity link function is used. When predicting probabilities of jumping, pup contact, or pup retrieval, *y* (*k*) denotes whether a certain event occurred during trial, *k* and a Binomial distribution with a logit link function is used. When predicting probabilities of behavioral statuses, *pup* (*k*), and *height* (*k*) are z-scored, and *y* (*k*) denotes the probability of being in a passive status (status 0) or active statuses (statuses 1, 2) in trial, *k* calculated over the period from trial start to landing, using 1-second time bins, and a binomial distribution with a logit link function is used. To analyze the effects of optogenetic stimulation, light+ and light-data were analyzed separately (Fig. 2b, e, f, h; Extended Data Fig. 2a-b, f).

### Calcium imaging and data preprocessing

Miniscope with a customized protective cap was connected to a rotary joint (Spinner, BN835106) and the Miniscope V4 data acquisition system. Calcium imaging data were collected at 10 Hz (30 Hz for one mouse #S2) and saved in TIFF format (.avi for mouse #S2) by the Bonsai software. Recordings were paused during inter-trial intervals (typically less than 5 minutes). Data recorded in the same session were pooled for analysis. The preprocessing pipeline consisted of eight steps to enhance signal quality and prepare data for subsequent analysis:

(1) **Glow Removal**: For each pixel, the minimum fluorescence value across the session was subtracted from all frames to mitigate glow effects caused by vignetting.
**Noise Reduction**: A spatial median filter with a 5 × 5 window was applied to reduce sensor-induced noise.
(2) **Background Subtraction**: A morphological opening operation was employed to estimate the background signal. The structuring element used was a circular kernel with a radius of 15 pixels, approximating the size of neurons in the imaging field. This operation first erodes and then dilates the image, effectively removing objects smaller than the structuring element. The background was estimated from the output of this operation, and subtracting it from the image yielded a clean foreground signal^50^.
(3) **Motion Correction**: The NoRMCorre algorithm with default parameters^51^ was used to correct motion artifacts. Briefly, NoRMCorre performs fast non-rigid motion correction by dividing the field of view (FOV) into overlapping spatial blocks and aligning each block to a dynamically updated template with subpixel precision. This approach approximates non-rigid motion as a series of piecewise rigid translations, allowing for the generation of a smooth motion field for each frame.
(4) **ROI Selection**: A maximum intensity projection was generated for each pixel across the session to facilitate manual selection of regions of interest (ROIs) corresponding to individual cells. Calcium signal data, denoted as *C*_*i*,1_ (*t*) were extracted for each cell *i* in time *t* based on these ROIs, where *t* represents time within the session.
(5) **Filtering**: Calcium signals from each trial were independently band-pass filtered (1/90–2.5 Hz) to eliminate baseline drift and high-frequency noise. To mitigate boundary effects, the first and last 200 data points of each trial were mirrored and appended as padding prior to filtering. Following filtering, the padding was removed, resulting in *C*_*i*,2_ (*t*) . For trials still exhibiting boundary artifacts, baseline correction was applied using a biexponential fitting model:

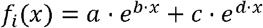

where *x* represents time within the trial. By aligning *f*_*i*_(*x*) with the corresponding time point of the session, *f*_*i*_(*t*) was obtained, represents the signal baseline for cell *i* at time *t* The corrected calcium signal was computed as:

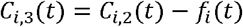
(6) **Gain and Baseline Adjustment**: Due to non-continuous recordings within a single session, signal gain and baseline differences could arise across trials. Gain correction was performed by selecting sparsely active cells, and calculating the standard deviation of the lower 65% of activity values in each trial. This standard deviation, denoted as *gain*_*i*,*k*_ represents the gain of cell *i* in trial *k*. A reference trial *r* was chosen, and the gain adjustment factor for each trial *g*_*k*_ was calculated as:

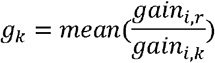 Here *i* only includes sparsely active cells chosen before. As cells recorded from the same trial tended to have a similar gain, the adjusted signal was computed as:

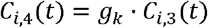

Baseline correction was then performed by calculating the 30th percentile of activity for each cell *i* in trial *k*, denoted as *b*_*i*,*k*_. The final adjusted signal was:

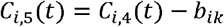

where each time point *t* was associated with its corresponding trial *k*.
(8) **Spike Inference**: Spike inference was performed using the OASIS algorithm^43^ with automatic parameter optimization. The algorithm models calcium signal dynamics using an autoregressive process and employs deconvolution to infer the spike sequence. Processed signals were then normalized via z-score transformation for further analysis, denoted as *s*_*i*_ (*t*).

### Analysis of cells responding to behavioral events

In Figure 3d, we calculated the average neuronal activity of each cell during the two seconds before and after the RA offset, JA offset, landing moment, and RT offset, and displayed the results. In panel (1), neurons were sorted based on the average neuronal activity during RA episode; in panel (2), neurons were sorted according to the average neuronal activity within a 1-second window after LA; and in panel (3), neurons were sorted based on the average neuronal activity during RT episode.

To determine whether a cell responds to a specific behavior, we used the following procedure. Taking risk assessment (RA) as an example, we calculated the average neuronal activity of each cell during each RA episode, with each episode contributing one data point to *Set* 1. For comparison, we considered the cell’s baseline activity at status 0. Baseline activity was smoothed using non-overlapping 2-second windows, and was denoted as *Set* 2. A two-sample t-test was performed to compare *Set* 1 and *Set* 2. If a cell’s average activity during RA episodes was significantly higher than the baseline, the cell was considered activated during RA. Conversely, if the activity was significantly lower, the cell was considered inhibited. Cells showing no significant difference were considered non-responsive to RA. The same approach was applied to identify cell responses to jumping attempt (JA) and retrieval (RT). For instantaneous events, such as wall down (WD) or landing (LA), specific time windows were analyzed to capture behavior-related neuronal activity: 1-second windows before WD and 1-second windows after LA. Extened Data Figure 4a illustrates the proportion of recorded cells responding to different behavioral events.

To test whether the fraction of cells responding to multiple events is significantly different from chance, we proposed a null model, in which neuronal response to each behavior is independent. We first calculated the proportion of activated cells for each event as the marginal probability, and estimated the expected proportion of cells for all combinatorial activation patterns. A binomial test was then performed to evaluate whether the observed number of activated cells for each activation pattern significantly deviated from the expected number. The significance levels were adjusted using the Bonferroni method.

Figure 3e illustrates the observed number of activated cells for each activation pattern, while Extended Data Figure 4b shows the deviance from chance, defined as:

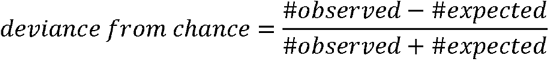

### Analysis of neuronal activity correlation across behaviors

We analyzed the activity of 641 neurons during five behaviors: IP, WD, RA, JA, RT, and LA. For IP, RA, JA, and RT, the analysis included all time points during which the mice exhibited the corresponding behavior. For WD, we analyzed the 1-second window before WD; for LA, we analyzed the 1-second window after LA. We computed the average neuronal activity across all time points associated with each behavior, and generated a 641 – by – 6 matrix. Pairwise Pearson correlations were next calculated to quantify population activity similarity across behaviors.

### Analysis of neural responses in different conditions

The heatmaps presented in Figure 4 and Extended Data Figure 4 show the average neural activity of activated individual cells aligned with the onset or offset of specific behavioral events (e.g., risk assessment, jumping attempt, and retrieval) after sorting. The related analysis was performed as follows, using Figure 4b as an example: (1) Cell selection. For cell *i*, total time bins were randomly divided into two equal subsets. Neural activity in subsets 1 and 2 was then calculated, and denoted as *s*_*i*1_ and *s*_*i*2_. To evaluate differences in neural activity between pup+ and pup-conditions, we focused on cells identified as significantly activated during the RA period in *s*_*i*1_. Selected cells were sorted based on average neural activity during the risk assessment (RA) period using the data from *s*_*i*1_. (2) Heatmap analysis. For selected cells, we further divided *s*_*i*2_ into two subsets based on the presence (*s*_*i*2+_) or absence (*s*_*i*2−_) of pups, and aligned its activity with the offset of RA (±2 s), separately for *s*_*i*2+_ and *s*_*i*2−_, using the cell order determined in the previous sorting step. (3) Statistical analysis. The mean neural activity during the RA period was calculated for each selected cell in *s*_i2+_ and *s*_i2−_, and a paired Wilcoxon signed-rank test was conducted. The procedures for the remaining panels in Figure 4 and Extended Data Figure 4 are similar to that used for Figure 4b, except that the event of interest was replaced by jumping attempt (JA) (Fig. 4c, e, g-i) or retrieval (RA) (Extended Data Fig. 4h, i), or aligned cells activity with the onset (± 2 s) of events (Extended Data Fig.4d, f, g, h and i). In Figure 4d, e, f g and Extended Data Figure 4h, i, the *s*_*i*2_ subset was divided based on the height of the platform during the recording. A generalized linear model (GLM) was used to predict average neural activity using heights, and the p-value of the height coefficient was used to determine statistical significance. In Figure 4h, the *s*_*i*2_ subsets were divided based on the outcome of the jumping attempt: retreat or landing. In Figure 4i, the time window of interest was 2 seconds after the JA onset. In Extended Data Figure 4i, the time window of interest was 2 seconds after the RT onset.

### Analysis of representational geometry

#### (1) Generative GLM models

Lasso GLM regression was used to model the generative processes underlying neuronal activity and behaviors (GLM1, GLM2, GLM6), with L1 norm. GLM1 predicted neuronal activity for each cell using height and presence of pups as predictors to assess how sensory inputs influence neuronal activity, using a normal distribution and an identity link function:

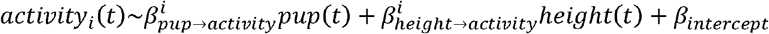

. GLM2 predicted behaviors using neuronal activity, height and presence of pups, using a binomial distribution and a logit link function:

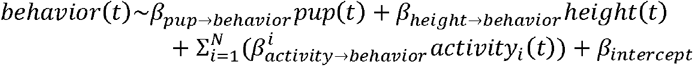

. GLM6 predicted neuronal activity for each cell using height and pup as predictors, restricted to time windows during which the mouse was immobile, using a normal distribution and an identity link function. This aimed to explore how sensory inputs influence neuronal activity before motor execution:

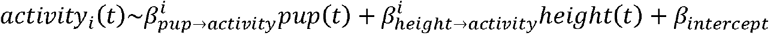

For each fitting, data recorded in the same session were pooled for analysis, and only the time period from trial onset to mouse landing for each trial was included. The variables include: (1) *activity*_*i*_ (*t*) which represents neuronal activity of cell *i* at time *t*, smoothed using a 3-second sliding window with 1.5-second overlaps; (2) *pup* (*t*), which represents whether a pup was placed in the trial at time *t*, and is one-hot encoded and z-score standardized; (3) *height* (*t*), which represents z-score standardized platform height of the trial at time *t* ; (4) *behaviour* (*t*), which is 1 when the mouse is engaged in risk assessment or a jumping attempt in more than 50% frames at time *t*, and 0 otherwise; (5) *β*_*intercept*_, which represents the intercept of each model; and (6) *β*_*x* →*y*_, which represents the coefficient of predictor *x* to predict outcome *y*, with only one valueper session, or 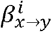, which represents the coefficient of predictor *x* to predict *y* specific to cell *i*, as shown in Figure 5b and 5c.

All models were trained using five-fold cross-validation. Cells with non-zero coefficients for pup or height in GLM1 were classified as sensory cells, while cells with non-zero coefficients for neuronal activity in GLM2 were classified as motor cells (Fig. 5g). Cells with both significant sensory and motor coefficients were used for analyses in Figure 5c and 5o. Cells with non-zero coefficients for pup, height, or neuronal activity were included in the analyses in Figure 5h, and the coefficients of these predictors were interpreted as pup effect, height effect, or motor effect, respectively.

#### (2) Low-dimensional visualization of representational geometry

To visualize representational geometry in high-dimensional space, we employed a GLM-QR method for dimensionality reduction. we first computed GLM1 coefficients as two *N* × 1 column vectors:

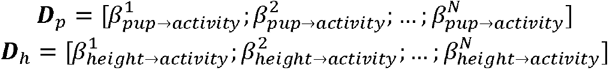

, where *N* is the number of recorded neurons. They represent pup and height coding directions. Similarly, we computed GLM2 coefficients:

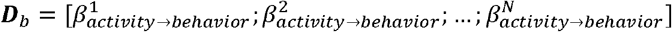

which represent the motivation-coding direction. Next, QR decomposition was applied to orthogonalize these coding directions:

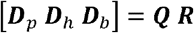

, where ***Q*** is an orthogonal matrix with orthonormal column vectors, satisfying ***Q***^*T*^ ***Q*** = ***I***, and ***R*** is an upper triangular matrix. The product ***Q***(;, 1: 3)^***T***^ · [***D***_*p*_ ***D***_*h*_ ***D***_b_] yields a 3 × 3matrix, where each column represents the projections of ***D***_*p*_ ***D***_*h*_ and ***D***_b_ onto the first three orthogonal subspaces defined by ***Q***, as shown in Figure 5d. Figure 5e and 5p display the cosine of the angles between the three coding directions computed for each session.

#### (3) Unique contribution model

A unique contribution model was employed to quantify the proportion of variance in neuronal activity uniquely attributable to motor or sensory inputs. The analysis was conducted on a per-cell basis using the following lasso generalized linear models with L1 norm (GLM3, GLM4, GLM5). GLM3 predicted neuronal activity using height, presence of pups, and behaviors as predictors, using a normal distribution and an identity link function:

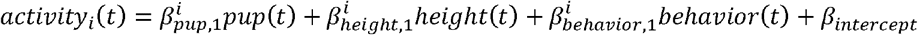

. Its model performance was reported as 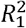. GLM4 predicted neuronal activity using height and presence of pups as predictors, using a normal distribution and an identity link function:

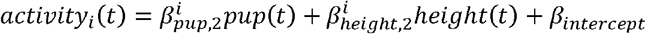

. Its model performance was reported as 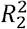. GLM5 predicted neuronal activity using behaviors as a predictor:

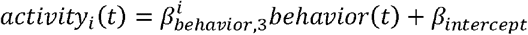

. Its model performance was reported as 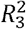. For each fitting, data recorded on the same session were pooled for analysis, and only the time period from trial onset to mouse landing for each trial was included. All models were trained using five-fold cross-validation. The variables here are the same as in the generative model described above. Next, the unique contributions of sensory and motor variables were calculated for each cell as 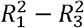 and 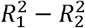, respectively

To assess chance levels for unique contributions, we performed 100 iterations of random shuffling. In motor shuffled model, we used height, presence of pups and shuffled behaviors as predictors to predict neuronal activity. Model performance was reported as 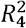. The difference 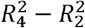 was used to estimate the chance level for motor unique contribution. In sensory shuffled model, we used behaviors, shuffled height, and shuffled presence of pups as predictors to predict neuronal activity. Performance was measured as 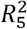. The difference 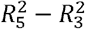 was used to estimate the chance level for sensory unique contribution. After 100 iterations, 100 shuffled motor and sensory unique contributions were calculated to determine chance levels. A cell was considered a significant motor or sensory unique cell if the corresponding unique contribution exceeded the upper bound of the corresponding 95% confidence interval (Fig. 5k, Extended Data Fig. 5b).

To assess whether a cell is sensory or motor biased, the preference index is defined as:

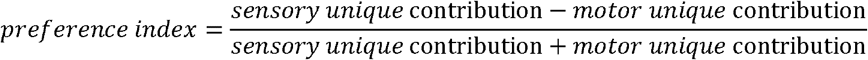

. When the preference index of a cell approaches 1, it indicates that the cell predominantly processes sensory information. Conversely, when it approaches -1, it suggests that the cell directly controls motor outputs.

### Dynamic models

To understand neural dynamics, we first sought to identify a motivation-coding direction in the activity space. Inspired by a previous study^4^, we derived a motivation axis and computed the projection of recorded population neural activity onto this axis, referred to as <u>A</u>ctivity in <u>M</u>otivation <u>D</u>imension (AMD):

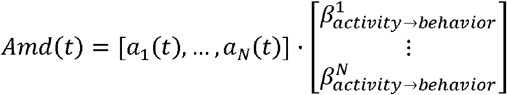

where *a*_*i*_(*t*) represents the activity of neuron *i* at time *t*, with neuronal spike data smoothed using a 1-second non-overlapping sliding window. 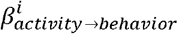 corresponds to the coefficients derived from the GLM2 model described earlier. *Amd* (*t*)is a scalar value that quantifies the mouse’s motivation at time *t*.

To investigate dynamics in motivation-coding dimension, we proposed five possible models and performed model comparison. For each model, data from all trials on a given session, spanning from trial start to mouse landing, were concatenated for parameter estimation.

#### (1) Gaussian model

The Gaussian Model assumes that the *Amd* data from a single session follow a Gaussian distribution with mean *µ* and standard deviation *σ*, and each data point is independent. The parameters were estimated as follows:

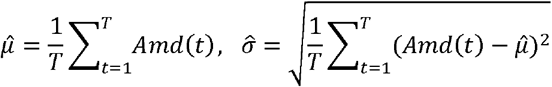

, where *T* denotes the total number of data points within the session. The negative log-likelihood (NLL) for this model is computed as:

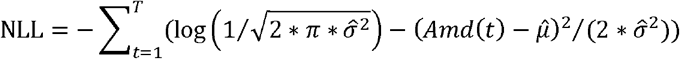

which quantifies the performance of the Gaussian model to the data.

#### (2) Autoregression model (AR)

The AR model assumes that *Amd* is generated through a first-order autoregressive process, where the value at each time point depends only on the value at the previous time point. The model is defined as follows:

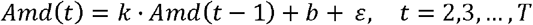

where *T* is the total number of data points within the session, is the autoregressive *k* is the autoregressive coefficient, *b* is the intercept, and *ε* represents the residuals, assumed to be gaussian noises with standard deviation *σ*.

This can be formulated as a generalized linear model (GLM), where each pair (*Amd* (*t* − 1), *Amd* (*t*)) represents a predictor-response data point. The parameters *k, b* and *σ* were estimated by solving the GLM.

The negative log-likelihood (NLL) for the AR Model is expressed as:

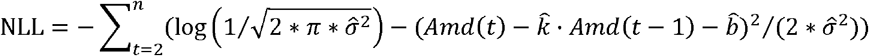

#### (3) Gaussian mixture model (GMM)

The Gaussian Mixture Model (GMM) assumes that *Amd* is generated from a mixture of two Gaussian distributions, corresponding to two latent states: state 1 and state 2. The probability density is defined as:

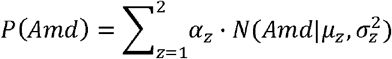

where *α*_z_ is the mixing weight, representing the probability of state *z*, and satisfying *α*_1_ + *α*_2_ = 1, and *N* denotes a Gaussian probability density distribution. The model parameters were estimated using the Expectation-Maximization (EM) algorithm, iterated for 200 cycles^52^.

##### E-step

Calculate the posterior probability *P*(*z*(*t*) | *Amd (t))*, where *z*(*t*) represents the state at time *t*:

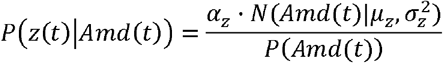

##### M-step

Update the model parameters

1. Mixing weights:

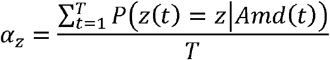

2. Means:

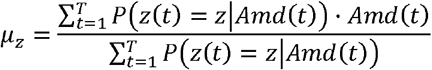

3. Standard deviations:

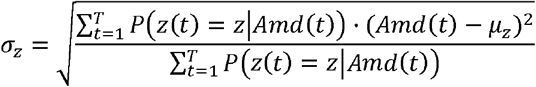

Negative log-likelihood (NLL):

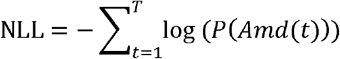

#### (4) Hidden Markov model - Gaussian model (HMM - Gaussian)

The HMM-Gaussian assumes that the observations (*Amd*) are dependent on a latent state characterized by a Markov process, which is not observable and needs to be inferred. The latent Markov process is characterized by stochastic switches between a resting state (state 1) and an active state (state 2), and the data in each state follow a Gaussian distribution. The transition probabilities are defined by the matrix *π*:

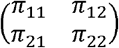

where *π*_*ij*_ = *P*(*z*(*t*) = *j* ∣ *z* (*t* − 1) = *i*).Gaussian distribution is defined as 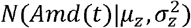. Parameters were estimated using the EM algorithm.

##### E-step

1. Forward Probability (Alpha): *α* (*t*,*z*) is defined as *α* (*t*,*z*) = *P* (*Amd*_1:*t*_, *z*_*t*_ = *z*), and calculated with:

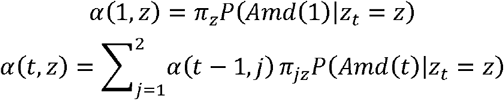
2. Backward Probability (Beta): *β*(*t*,*z*) is defined as *β*(*t*,*z*) = *P* (*Amd*_(*t*+1):*T*_ ∣ *z*_*t*_ = *z*),and calculated with:

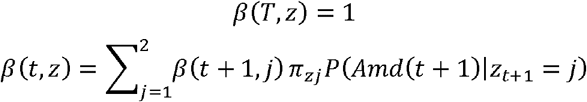
3. Posterior probabilities *P* (*z*(*t*) ∣ *Amd* (1:T)):

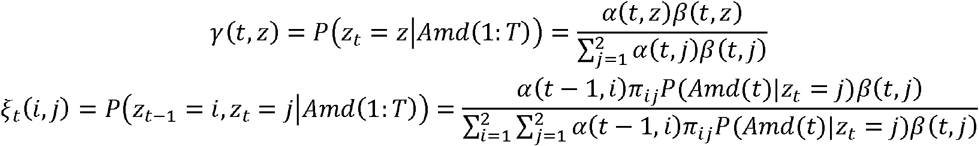

##### M-step

1. Update the transition matrix:

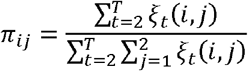
2. Update Gaussian means and variances:

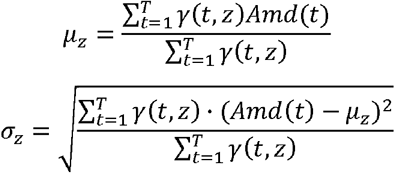

Negative log-likelihood (NLL):

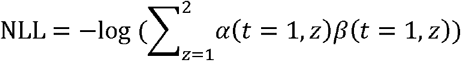

#### (5) Hidden Markov model - Autoregression model (HMM - AR)

The HMM-AR model also assumes that the observations (*Amd*) are dependent on a latent Markov process (state). Similar to HMM-Gaussian, the latent Markov process is characterized by stochastic switches between a resting state (state 1) and an active state (state 2). However, in HMM-AR, the data in each state are generated through a first-order autoregressive process. The state transition probabilities are represented by a matrix *π*:

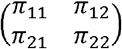

where *π*_*ij*_ = *P* (*z*(*t*) = *j* ∣*z* (*t* −1) *= i*. The observed data *Amd* are assumed to follow an autoregressive process. Specifically, the activity at the current time point depends solely on the activity at the previous time point *Amd*_*t*-1_ and the state at the current time *z*(*t*):

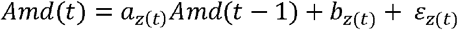

where 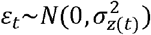 is a Gaussian noise. The parameters were estimated using the EM algorithm.

##### E-step

1. Forward Probability (Alpha): *α* (*t*,*z*) is defined as *α* (*t*,*z*) = *P* (*Amd*_1:*t*_, *z*_*t*_ = *z*), and calculated with:

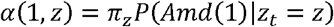

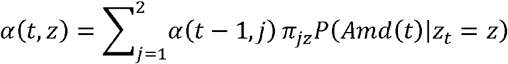
2. Backward Probability (Beta): *β*(*t*,*z*) is defined as *β*(*t*,*z*) = *P* (*Amd*_(*t*+1):*T*_ ∣ *z*_*t*_ = *z*),and calculated with:

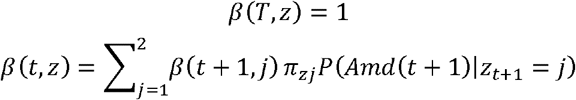
3. Posterior probabilities *P* (*z*(*t*) ∣ *Amd* (1:T)):

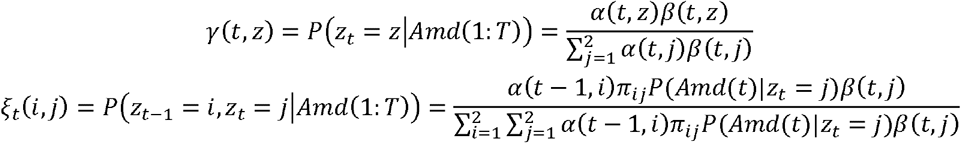

##### M-step

1. Update the transition matrix:

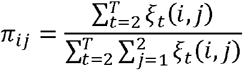
2. Update AR coefficients: Estimate the coefficients *a* and *b* using weighted least squares:

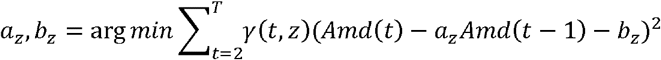
3. Update variance:

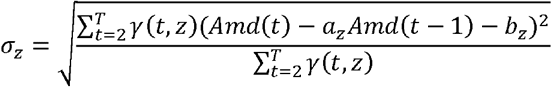 Negative log-likelihood (NLL):

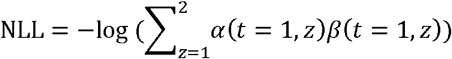

For the three models involving the Expectation-Maximization (EM) algorithm (GMM, HMM-Gaussian, and HMM-AR), the algorithm was iterated 200 times to ensure convergence. To minimize the risk of being trapped in local optima, the EM procedure was repeated ten times with random initializations of the parameters. The parameter set corresponding to the highest likelihood was selected for subsequent analyses. After parameter estimation, the most likely sequence of latent states at each time point was inferred by maximizing *P*(*z*(*t*) | *Amd*) per time point.

To determine the most suitable model for each session’s data, we employed three evaluation criteria: Akaike Information Criterion (AIC), Bayesian Information Criterion (BIC), and cross-validation. AIC and BIC are defined as:

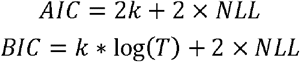

where *k* denotes the number of model parameters (Gaussian: 2, AR: 3, GMM: 5, HMM-Gaussian: 6, HMM-AR: 8), and *T* is the total number of data points analyzed for the given session. Both AIC and BIC prioritize models with smaller values, with lower scores indicating better model performance. For cross-validation, data of a session were divided into non-overlapping 60-second epochs. A four-fold cross-validation approach was implemented as follows: three-quarters of the epochs were randomly selected for model training, and the remaining epochs were used to compute the negative log-likelihood (NLL) based on the estimated parameters. This process was repeated four times, and the average NLL across the four folds was taken as the cross-validated NLL of the session. Similar to AIC and BIC, lower cross-validated NLL values indicate better model performance.

To explore the influence of pup and platform height on neural dynamics, we categorized data from each session into eight conditions based on pup and height variables. For each condition *u* on session *d*, the mostly likely latent state sequence *z*_*d*,*u*_ (*t*) was inferred using the HMM-AR model fitted on all trials^53^. Then transition probabilities and autoregressive parameters were calculated for each condition separately:

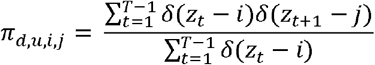

where 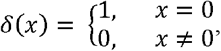 and

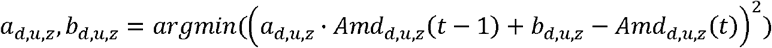

. Once the parameters were determined, the following variables were computed for each condition in each session. (1) Probability of staying: *P*1_*d*,*u*_ = *π*_*d*,*u*,1,1_, and *P*2_*d*,*u*_ = *π*_*d*,*u*,2,2_, representing probability of staying in the previous state (2) Relaxation time: *tau*_*d*,*u*_(*z*) = 1/ (1-*a*_*d*,*u*,*z*_), representing the growth and decay time constant in the autoregressive process in a given state. (3) Asymptotic activity: *M*_*d*,*u*_(*z*) = *b*_*d*,*u*,*z*_/ (1-*a*_*d*,*u*,*z*_), representing the steady-state neural activity.To assess the effects of pup and height on these parameters, we pooled data across all sessions and fitted the parameters using a generalized linear model (GLM):

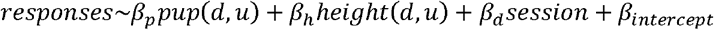

Predictors include: (1) *pup*(*d*,*u*): The presence of pups for condition *u* on session *d* (one-hot encoded and z-score standardized). (2) *height* (*d*,*u*): The platform height for condition *u* on session *d* (z-score standardized). (3) *day*: The recording session index (one-hot encoded) to incorporate variations across sessions and individuals. Responses include: *P*1 *d*,*u, P*2 *d*,*u, tau*_*d*,*u*_(l), *tau*_*d*,*u*_(2), *M*_*d*,*u*_(l), *M*_*d*_,_*u*_(2). The GLM results are presented in Figure 6j.

### Autocorrelation analysis

To demonstrate that the ramping activity and hence autocorrelation observed in the motivation dimension is not simply an artifact of imperfect deconvolution of slow calcium signals, we randomly shuffled the motivation dimension vector in GLM2, and project neural activity onto the shuffled dimension:

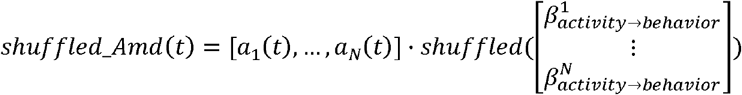

. This shuffling procedure was repeated 200 times. We computed the autocorrelation of *shuffled_Amd* for each iteration and *Amd*(Extended Data Fig. 6).

## Data availability

Data of the study are available from the corresponding authors upon reasonable request.

## Code availability

Codes of the study are available from the corresponding authors upon reasonable request.

## Acknowledgements

We thank S. Shea and S. Musall for critical feedback on the manuscript; X. Zhao, X. Guo, Y. Fu and T. Meng for equipment support; UCLA Miniscope Google group for helpful advice; and all members of the Huang lab for helpful discussion. This study was supported by Science and Technology Innovation 2030--Brain Science and Brain-inspired Intelligence Project of China 2022ZD0206900 (to L.H.); National Natural Science Foundation of China 32271143 (to L.H.) and 32300839 (to Y.X.); China Postdoctoral Science Foundation 2021M703417 and 2023T160669 (to Y.X.).

## Author contributions

Y.X. and L.H. conceptualized the study, designed experiments, analyzed data and wrote the manuscript. L.H. supervised the project. Y.X. performed most of the experiments and prepared figures. Y.L. analyzed data, performed computational modeling, prepared related figures and wrote the manuscript. X.D. assisted in behavior experiments. All authors revised the manuscript.

## Supplementary figure legend

**Extended Data Figure 1.**
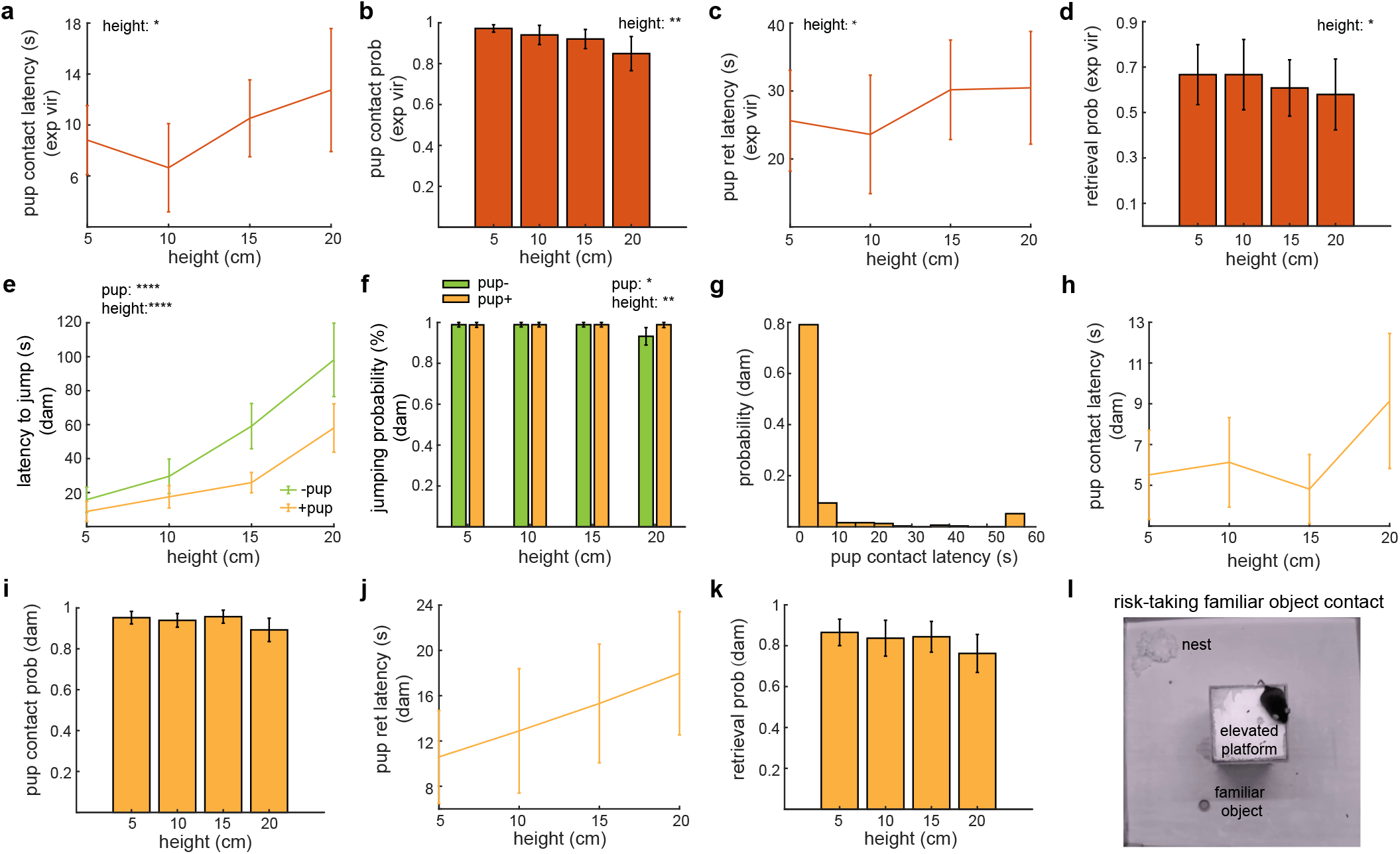
Pup-oriented motivation drives mice to overcome fear of height in risk-taking maternal behaviors. **a-b**. Pup contact latency and probability in experienced virgin mice. **c-d**. Latency and probability of retrieving the first pup in experienced virgin mice. **e-f**. Latency to jump and jumping probability in dams. **g**. Distribution of pup contact latency in dams. **h-k**. Same as **a-d**, but in dams. **l**. Illustration of the risk-taking familiar object contact behavior paradigm. Data are mean ± s.e.m. *p<0.05, ****p<0.0001, GLM. n=7 mice for the experienced virgin mice (**a-d**) and n=7 mice for the dams (**e-k**).

**Extended Data Figure 2.**
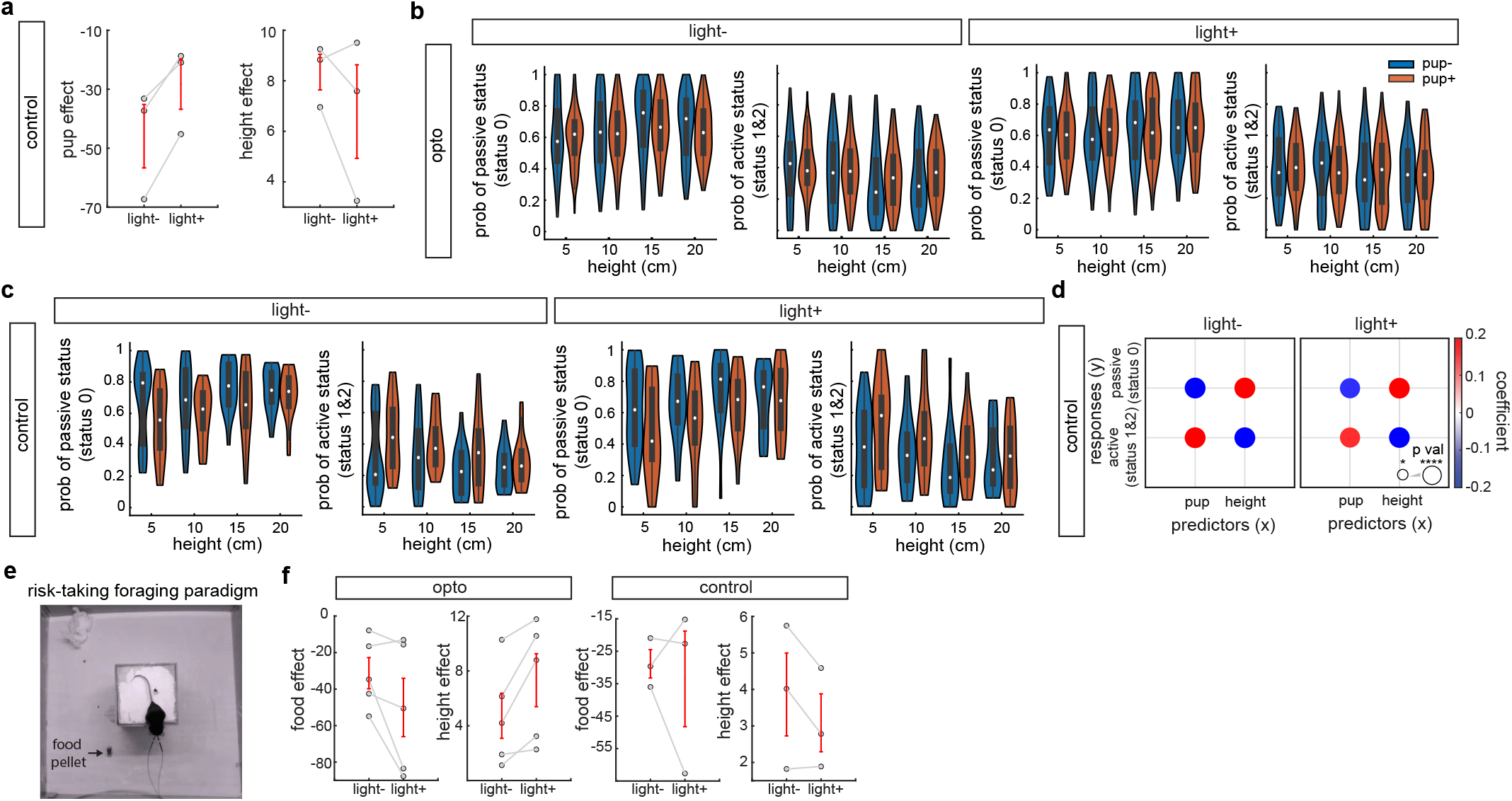
Optogenetic inhibition results in reduced pup-oriented motivation in risk-taking maternal behaviors. **a**. The pup effect (left) and the height effect (right) in light-versus light+ conditions for the control group. **b-c**. Probability of the passive status (status 0) and active status (status 1 and 2) in light- and light+ conditions in the opto (**b**) and control (**c**) groups. The white dots, boxes, and whiskers represent the median, interquartile range (IQR), and the range extending from 1.5×IQR below the lower quartile to 1.5×IQR above the upper quartile, with whiskers clipped to the data range respectively. **d**. Effects of pup and height on the probability of the passive and active status in the light-(left) and light+ (right) groups of the control group. Colors and sizes of circles effect represent weights (GLM coefficients) and significance levels (p value). **e**. Illustration of the risk-taking foraging behavior paradigm. **f**. Food and height effects in light-versus light+ conditions in the opto (left) and control (right) groups. Data are mean ± s.e.m. (**a, f**). n=13 mice (opto group-pup), n=3 mice (control group-pup) (**a-d**) n=5 mice (opto group food) and n=3 mice (control group -food) (**f**).

**Extended Data Figure 3.**
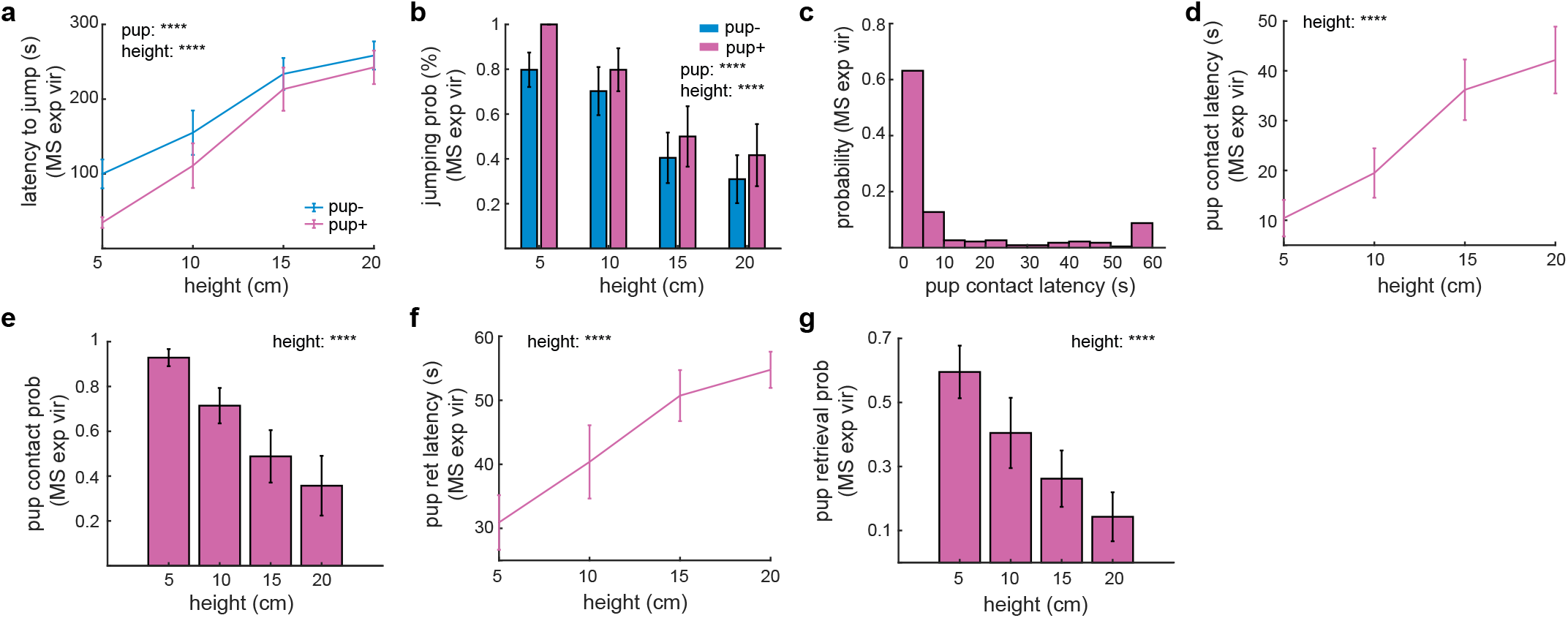
Experienced virgin females for miniscope recording exhibit maternal motivation in risk-taking maternal behaviors. **a-b**. Latency to jump and jumping probability. **c**. Distribution of pup contact latency. **d-e**. Pup contact latency and probability. **c-d**. Latency and probability of retrieving the first pup. Data are mean ± s.e.m. ****p<0.0001, GLM. n=7 experienced virgin mice for miniscope recording.

**Extended Data Figure 4.**
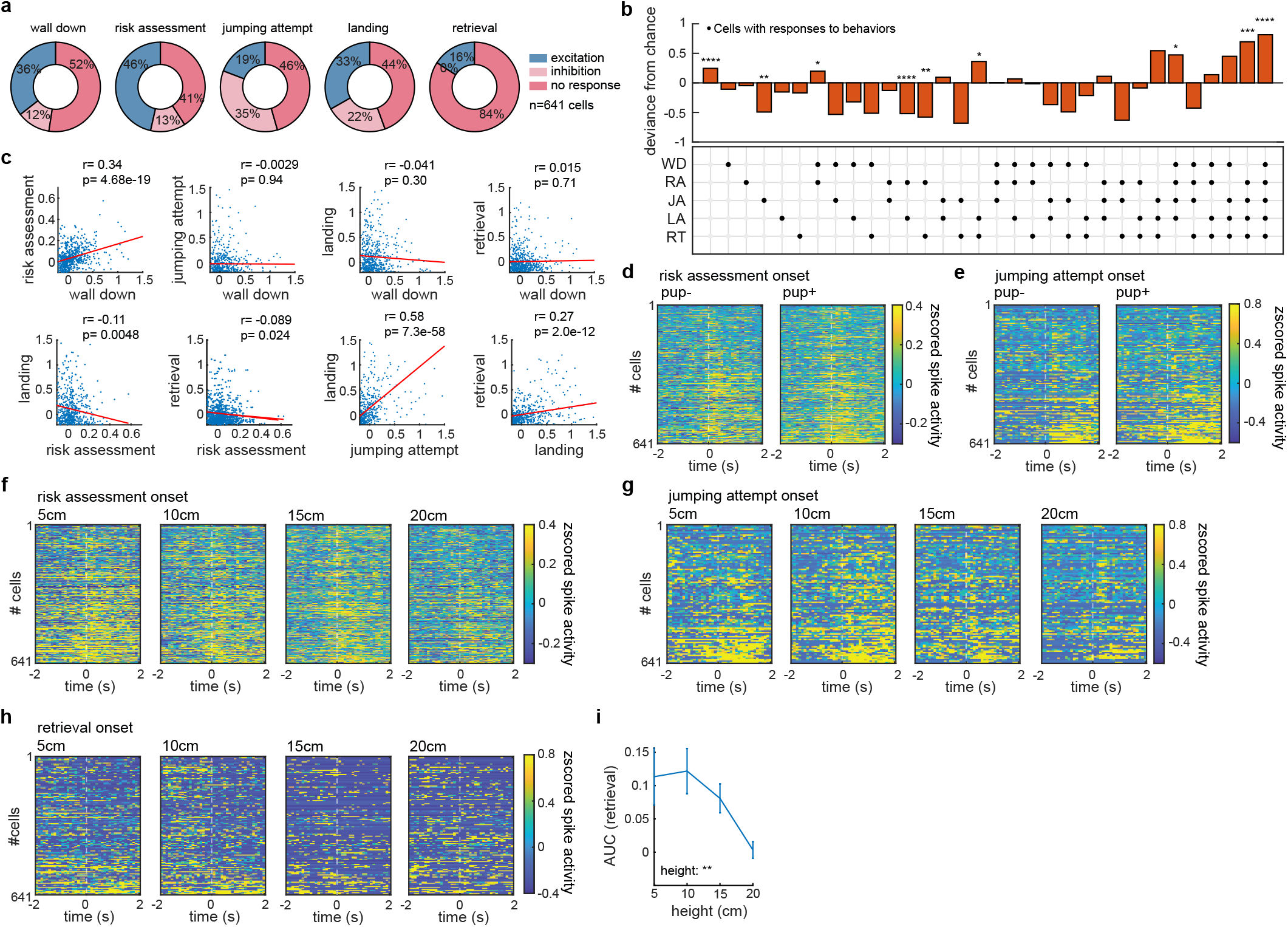
The PAG-projecting mPFC neurons encode maternal motivation for risk-taking maternal behaviors. **a**. Proportions of recorded neurons that were excited, inhibited or had no responses in various behaviors. **b**. Deviance of the number of activated cells in different behaviors from chance. See methods for statistical analysis. *, **, *** and **** denote Bonferroni corrected p values < 0.05, 0.01, 0.001 and 0.0001 respectively. **c**. Scatter plots showing average activity between pairs of behavioral events. Each dot represents an individual neuron. The x or y axis represents average activity during a certain behavior. Red lines represent the fitted lines. r, Pearson correlation coefficients. p, p values. **d**. Heatmaps of cross-validated z-scored spike activity aligned with the onset of risk assessment. Neurons are sorted by their activity during the entire event of risk assessment. **e**. Same as d, but aligned with the onset of jumping attempt. **f.-g**. Heatmaps of cross-validated z-scored spike activity aligned with the risk assessment onset (**f**) and the jumping attempt onset (**g**) at different heights. Neurons are sorted by their activity during the entire events. **h**. Heatmaps of cross-validated z-scored spike activity aligned with the onset of pup retrieval. Neurons are sorted by their activity in the 2 s windows after the pup retrieval onset. **i**. AUC of retrieval at different heights. Data are mean ± s.e.m. **p<0.01, GLM. See Supplementary Information for the number of cells for each panel. Cells were from 20 sessions in 7 mice.

**Extended Data Figure 5.**
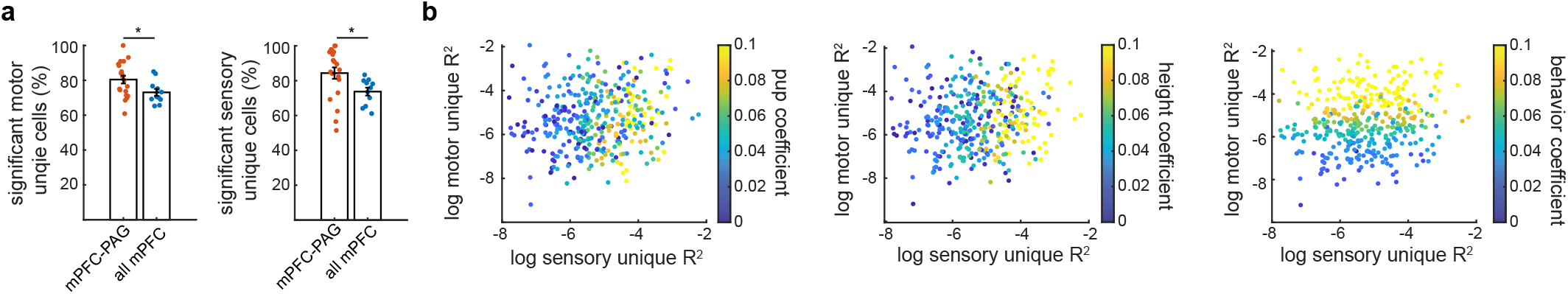
Representational geometry allows for integration of pup and height cues to drive pup-oriented risk-taking behaviors, related to Fig. 5. **a**. Percentage of neurons with significant unique sensory or motor contributions. Data are mean ± s.e.m. *p<0.05, two-sample t-test. **b**. Scatter plots of sensory unique R^2^ versus motor unique R^2^ color coded with *β*_*pup* → *activity*,_ *β*_*height*→ *activity*_. and *β*_*activity*.→ *behaviour*_. Only neurons with significant motor and sensory unique R^2^ in Fig. **5k** are shown. n=641 (**a**) and 384 (**b**) PAG-projecting mPFC cells from 20 sessions in 7 mice, and n=1160 pan mPFC cells (**b**) from 11 sessions in 3 mice (see methods).

**Extended Data Figure 6.**
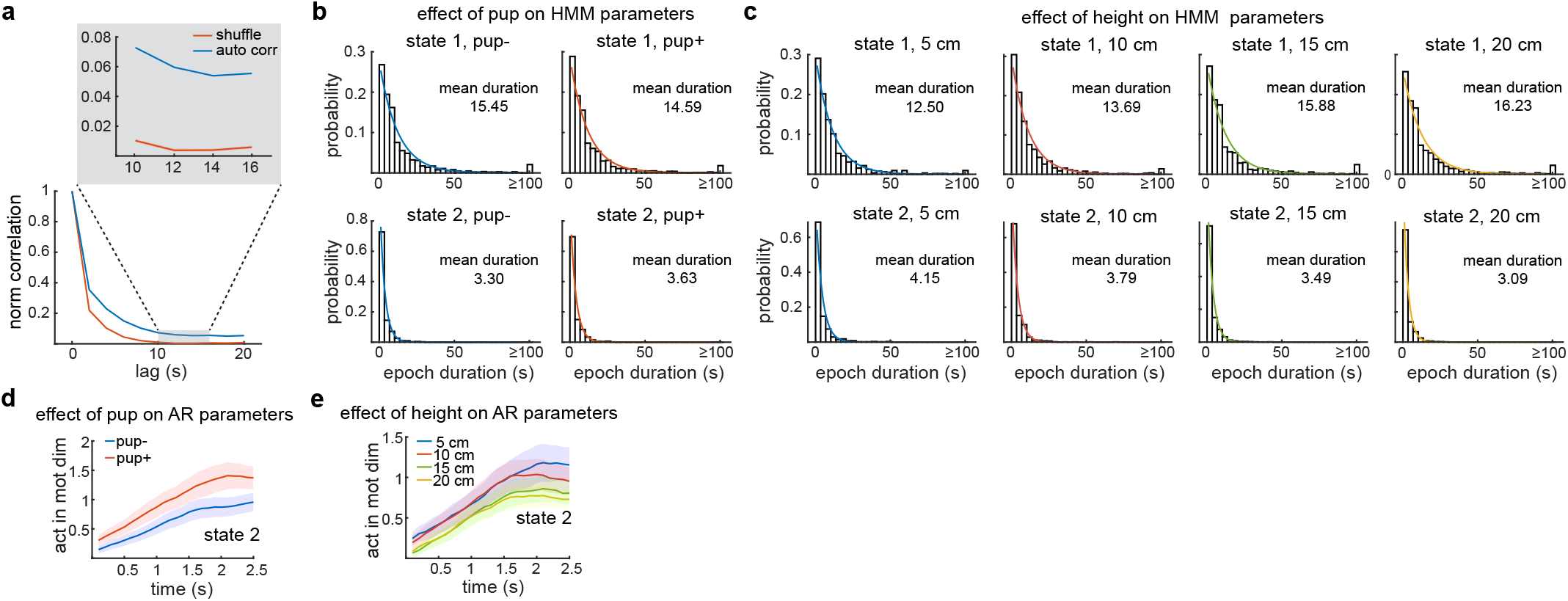
Pup and height cues oppositely modulate HMM and AR parameters. **a**. Normalized autocorrelation of spike activity in motivation-encoding dimension versus in random dimensions (shuffle). Neural activity in the motivation-encoding dimension displayed significantly higher autocorrelation. Shuffled data are presented as mean with 95% confidence intervals. **b-c**. Effects of pup and height on the distribution of epoch durations of state 1 and 2 for the HMM parameters. **d-e**. Effects of pups and height on the activity in the motivation dimension during state 2 in different conditions for the AR parameters. Data are mean ± s.e.m. (**d, e**). n=641 neurons from 20 sessions in 7 mice.

**Extended Data Figure 7.**
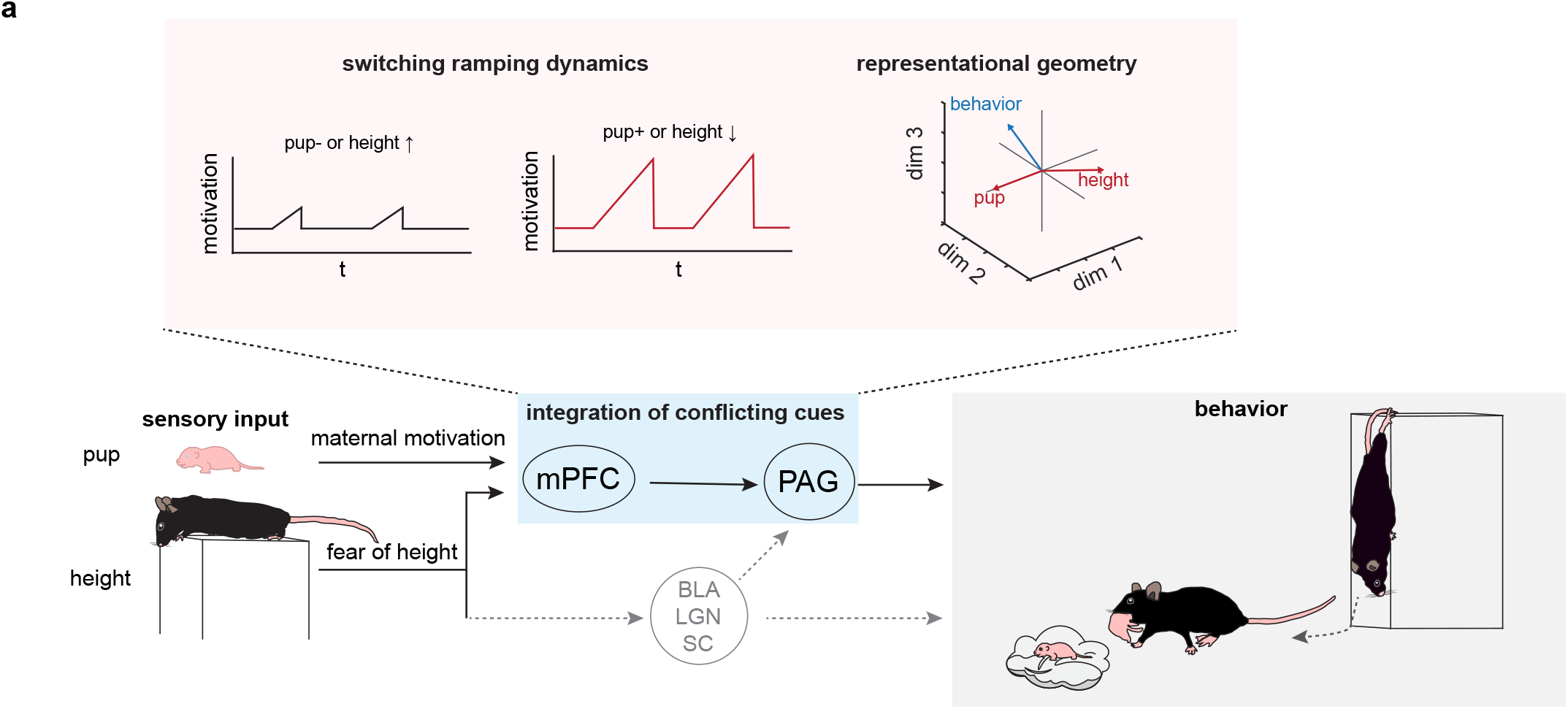
A model summarizing neural mechanisms of risk-taking maternal behaviors. **a**. Switching ramping dynamics and representational geometrics allow for integration of pup and height cues to dynamically modulate motivation, which give rises to risk-taking maternal behaviors.

